# Glucocorticoid intermittence coordinates rescue of energy and mass in aging-related sarcopenia through the myocyte-autonomous PGC1alpha-Lipin1 transactivation

**DOI:** 10.1101/2023.10.16.562573

**Authors:** Ashok Daniel Prabakaran, Kevin McFarland, Karen Miz, Hima Bindu Durumutla, Kevin Piczer, Fadoua El Abdellaoui Soussi, Hannah Latimer, Cole Werbrich, N. Scott Blair, Douglas P Millay, Brendan Prideaux, Brian N Finck, Mattia Quattrocelli

**Affiliations:** Molecular Cardiovascular Biology, Heart Institute, Cincinnati Children’s Hospital Medical Center and Dept. Pediatrics, University of Cincinnati College of Medicine, Cincinnati, OH, USA; Department of Neuroscience, Cell Biology, and Anatomy, University of Texas Medical Branch (UTMB), Galveston, TX, USA; Department of Medicine, Center for Human Nutrition, Washington University in St Louis, MO, USA

**Keywords:** Glucocorticoid steroids, muscle mitochondria, muscle mass, sarcopenia, inducible myocyte-specific PGC1alpha knockout, PGC1alpha isoforms, inducible myocyte-specific Lipin1 knockout

## Abstract

Sarcopenia burdens the elderly population through loss of muscle energy and mass, yet treatments to functionally rescue both parameters are missing. The glucocorticoid prednisone remodels muscle metabolism based on frequency of intake, but its mechanisms in sarcopenia are unknown. We found that once-weekly intermittent prednisone rescued muscle quality in aged 24-month-old mice to levels comparable to young 4-month-old mice. We discovered an age- and sex-independent glucocorticoid receptor transactivation program in muscle encompassing PGC1alpha and its co-factor Lipin1. Treatment coordinately improved mitochondrial abundance through isoform 1 and muscle mass through isoform 4 of the myocyte-specific PGC1alpha, which was required for the treatment-driven increase in carbon shuttling from glucose oxidation to amino acid biogenesis. We also probed the myocyte-specific Lipin1 as non-redundant factor coaxing PGC1alpha upregulation to the stimulation of both oxidative and anabolic capacities. Our study unveils an aging-resistant druggable program in myocytes to coordinately rescue energy and mass in sarcopenia.

## Introduction

One of the most consequential phenotypes of aging is the multifaceted decline in skeletal muscle function, which contributes to loss of mobility and affects lifestyle in the elderly population ^1^. Muscle aging is a particularly significant context to study the relationship between bioenergetics and mass remodeling. With aging, muscle loses mass (sarcopenia) and quality, i.e. intrinsic capacity of generating force ^2^. Sarcopenia and geriatric weakness correlate with impaired metabolic capacity to produce energy in muscle ^3,4^. However, the reciprocal regulations between metabolic capacity and mass remodeling in muscle aging remain largely unelucidated. Considering the ever-increasing aging of our population in the US and worldwide, it is imperative to identify pharmacological targets for balanced recovery of muscle quality and mass in advanced age.

While the role of peroxisome proliferator-activated receptor-gamma coactivator 1 alpha (PGC1alpha) in mitochondrial capacity ^5^ and overall mitochondrial protein quality^6^ is quite established, its effects on age-related sarcopenia and weakness are still debated with conflicting results. PGC1alpha impact on metabolism is balanced at the level of its splice variants^7^. Besides the canonical longer PGC1alpha-isoform 1 that regulates mitochondrial biogenesis and function, the shorter PGC1alpha-isoform 4 has been identified in mice and humans from an alternative transcription start site^8^ and is sufficient to increase muscle mass and strength in cachectic muscle^9^ and sarcopenia^10^. A previous study with caloric restriction reported correlations between PGC1alpha upregulation in aging muscle and activation of growth pathways in addition to mitochondrial function ^11^. Concordantly, studies in aging transgenic mice reported gain of muscle mass with constitutive PGC1alpha overexpression ^12^ and, conversely, loss of lean mass with constitutive muscle PGC1alpha knockout ^13^. However, another study with constitutive PGC1alpha overexpression versus knockout in muscle showed that PGC1alpha was dispensable for age-related sarcopenia ^14^. Another recent study showed that lifelong muscle PGC1α overexpression increased muscle mass in males but not females, and improved muscle fatigue at the expense of specific force ^15^. Thus, the role of myocyte-specific PGC1alpha in rescuing age-related sarcopenia and weakness remains unclear. This opens the question of whether additional factors balance the PGC1alpha action on energy and mass in muscle. Notably, the mechanisms coaxing mitochondrial metabolism to mass growth in the aging muscle are still largely unknown.

Lipin1 is a multi-functional protein that regulates muscle function and bioenergetics, and its ablation leads to muscle dysfunction and lipid accumulation in mice ^16^. Lipin1 acts in the cytosol as a phosphatidic acid phosphohydrolase ^17,18^ and in the nucleus as a regulator of gene transcription ^19^. In muscle, Lipin1 regulates many complex processes, including myofiber stability and regeneration ^20^, as well as autophagy/mitophagy ^21,22^. In hepatocytes, Lipin1 co-activates PGC1alpha through a direct protein-protein interaction ^19^, but this role of Lipin1 remains unexplored in muscle. More generally, the role of the myocyte-specific Lipin1 in muscle aging and energy-mass balance requires further investigation.

Glucocorticoid steroids are potent drugs that regulate both energy metabolism and mass. Traditionally, these drugs are considered for their immunomodulatory effects, as in genetic dystrophic myopathies ^23^. However, experiments with in vitro patient-derived myotubes clearly showed glucocorticoid-driven benefits in dystrophic muscle cells in the absence of immune cells ^24^. Furthermore, dosing frequency of glucocorticoid intake determines the benefits/risks ratio of these drugs with regards to metabolic balance. Chronic once-daily glucocorticoid intake promotes metabolic imbalance ^25^. Conversely, dosing intermittence shifts the glucocorticoid metabolic program from **pro-wasting**, i.e. atrophy and decreased bioenergetics with once-daily prednisone, to **pro-ergogenic**, i.e. increased bioenergetics and muscle mass with once-weekly prednisone in young adult mice, counteracting the muscle detriments induced by diet-induced obesity ^26^. In dystrophic patients, a recent pilot clinical trial reported positive trends in both lean mass and mobility with once-weekly prednisone ^27^. However, the relevance of glucocorticoid intermittence in the context of muscle aging is still unknown. In that regard, the myocyte-autonomous mechanisms of glucocorticoid drugs that could benefit the aging muscle are still undefined.

Here we report on the rejuvenating effects of intermittent prednisone on both bioenergetics and mass in the aging muscle of male and female mice. We interrogated transcriptomic and epigenomic datasets to identify activation of Lipin1-PGC1alpha axis. We used inducible myocyte-specific knockout models for PGC1alpha and Lipin1 to investigate requirement of these factors for the coordinated rescue of energy and mass in the absence of developmental or lifelong muscle adaptations to the manipulation of those genes. Moreover, we found that the PGC1alpha upregulation mediates the boost in amino acid biogenesis from oxidative intermediates, linking the bioenergetic and anabolic stimulations of treatment in muscle. Our study provides evidence and myocyte-specific mechanisms to challenge existing paradigms on glucocorticoid drugs with unexpected anti-sarcopenic effects.

## Results

### Intermittent once-weekly prednisone rejuvenates mitochondrial and mass properties of the aging muscle

Muscle aging is characterized by declines in both mitochondrial capacity and mass ^28^. Based on the initial positive effects we documented on both mitochondrial function and mass in young adult muscle of WT mice ^29^, we tested the extent to which an intermittent once-weekly prednisone treatment impacted muscle properties in the context of aging. We therefore treated aged WT mice at 24 months of age and the background-matched (*C57BL/6JN*) young adult controls at 4 months of age, all from the National Institute on Aging’s Division of Aging Biology mouse colony. Treatment was consistent with our prior report ^29^, i.e. once-weekly 1mg/kg prednisone i.p. at ZT0 for 12 weeks, controlled by the same schedule of vehicle administration. We treated males and females in parallel and data are reported as disaggregated by sex in **fig1** and **Suppl. Fig. 1**.

**Figure 1.**
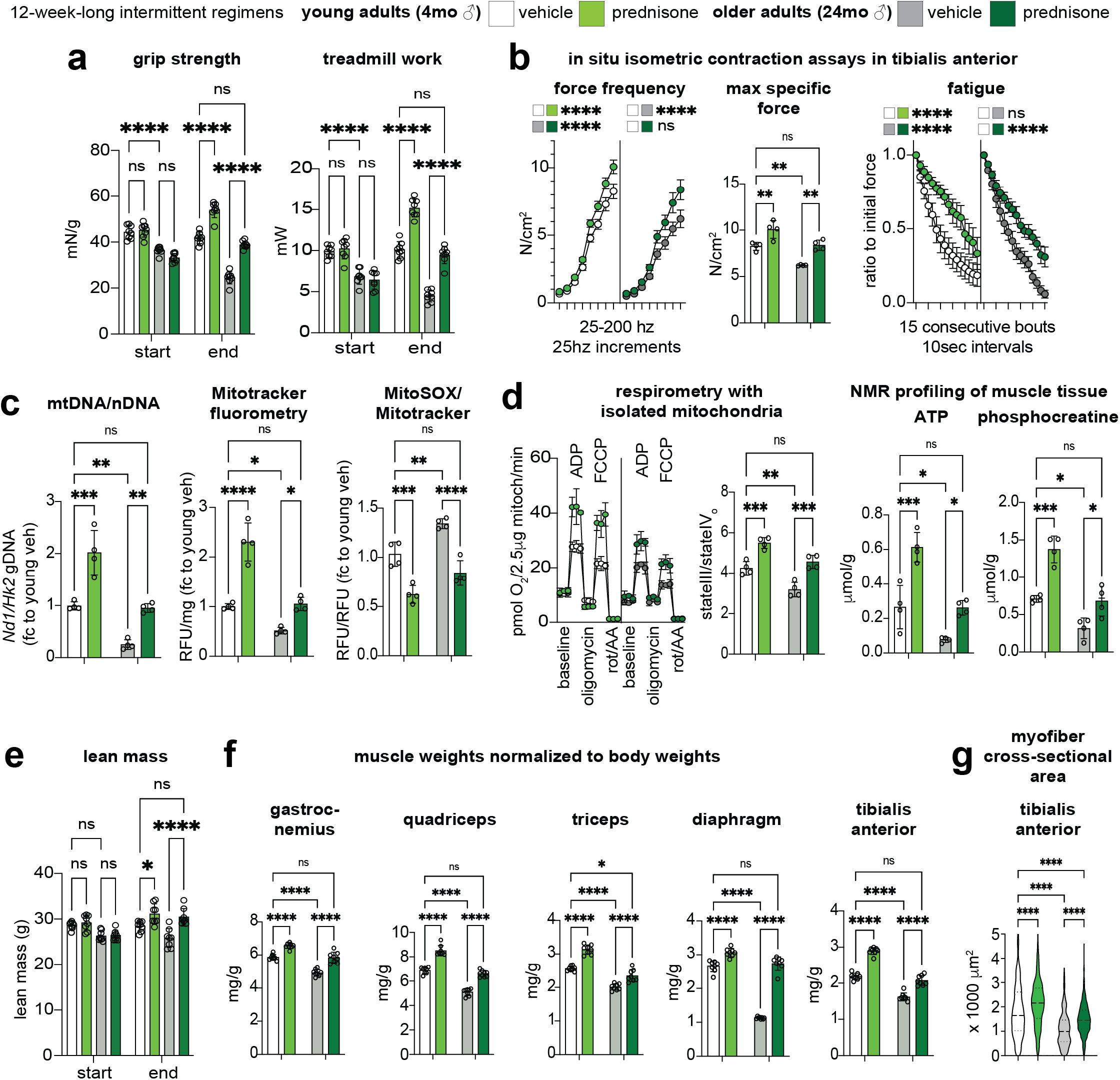
Intermittent once-weekly prednisone regimen rejuvenates mitochondrial and mass properties of the aging muscle. **(a)** Treatment improved strength and treadmill performance in background-matched male mice at young adult (4mo) and older adult (24mo) ages, improving the parameters of treated aged mice to levels comparable to the control (vehicle) young adult mice at endpoint. **(b)** Treatment rescued specific force in older mice to control young levels, while increasing resistance to repetitive tetanus fatigue to comparable extent at both ages. **(c-d)** Treatment improved mitochondrial abundance (mtDNA/nDNA, Mitotracker) and decreased ROS levels (MitoSOX) in aged muscle to young control-like levels. Analogous trends were observed with mitochondrial respiration levels and NMR-quantitated levels of ATP and phosphocreatine in quadriceps muscles. **(e-g)** In treated older mice, total lean mass increased to young control-like levels. This correlated with rescue of muscle weight/body weight ratios in older mice in locomotory (gastrocnemius, quadriceps, triceps) and respiratory (diaphragm) muscles. Tibialis anterior muscle analyses showed coupling of myofiber CSA trends with the changes in muscle mass. N=4-8/group; 2w ANOVA + Sidak: *, P<0.05; **, P<0.01; ***, P<0.001; ****, P<0.0001.

As parameters of overall strength and function, we quantitated grip strength and treadmill performance at baseline and after treatment (i.e. ∼27 months of age in older mice), and quantitated force production in situ in tibialis anterior muscles after treatment. At baseline, compared to young controls, older mice showed decreased strength and treadmill endurance. Compared to vehicle, treatment increased both parameters in older mice at endpoint. The values for treated older mice were not significantly different from the values exhibited by the control vehicle young mice at endpoint. As a validation, the treatment effect was recapitulated in the young mice **(fig1)**. At endpoint, we used isometric contraction assessments to profile force production through both force-frequency and fatigue assays. Treatment improved specific force in older muscle to levels like the ones shown by the control young muscle, while resistance to fatigue was improved by treatment to similar extents in both age groups **(fig1).**

We then analyzed mitochondrial properties consistently with our previous treatment study ^29^. We measured relative trends in mitochondrial abundance through mtDNA/nDNA (mitochondrial/nuclear DNA) qPCR quantitation and unbiased Mitotracker fluorometry in parallel in isolated myofibers from the flexor digitorum brevis muscles. Together with Mitotracker, MitoSOX fluorometry was used to quantitate ROS production. Treatment increased mitochondrial abundance in both age groups, rescuing the values of treated older muscles to young control-like levels. Conversely, ROS production was decreased by treatment in older muscle, suggesting functional coupling in the overall mitochondrial pool after treatment **(fig1)**. This was further elucidated in quadriceps muscle by respirometry curves with isolated mitochondria (fuel: pyruvate) and NMR-based quantitation of the energy-exchange molecules ATP and phosphocreatine. Treated aging muscle showed young control-like levels of ADP-fueled respiratory control ratio (state III/stateIV_o_, ^30^) and static levels of ATP and phosphocreatine **(fig1)**.

We then analyzed parameters of lean and muscle mass. Using echoMRI, we found that treatment increased overall lean mass in treated older mice to levels similar to the ones found in young control mice **(fig1)**. The trends in lean mass were matched by analogous trends in muscle/body weight ratios throughout the body, as shown by measures in four different locomotory muscles (gastrocnemius, quadriceps, triceps and tibialis anterior) and the respiratory muscle diaphragm **(fig1)**. The trends in muscle/body weight were further matched by analogous trends in myofiber cross-sectional area, as shown in the case of tibialis anterior **(fig1)**, further illustrating the treatment-driven rescue of aging muscle mass to young control-like levels. Discussed so far were treatment effects in male cohorts, but analogous trends were recorded in parallel for age- and background-matched female cohorts from the same colony **(Suppl. Fig. 1)**. Moreover, we did not record sizable treatment effects on top of the expected age-related shifts in relative myofiber type abundances in two locomotory muscles with mixed fiber typing, i.e. gastrocnemius and triceps, in both sexes **(Suppl. Fig 2)**.

Thus, according to the drug schedule and readout parameters tested here, intermittent prednisone “rejuvenated” both mitochondrial function and mass in the aging muscle, i.e. improved parameters in treated older muscles to young control-like levels.

### Treatment induces a muscle GR program increasing PGC1alpha-Lipin1 expression through aging

To gain insight in the mechanisms mediating the dual treatment effects on energy and mass in aging muscle, we profiled the epigenomic signal of the glucocorticoid receptor (GR) in all age/sex/treatment cohorts, in parallel with bulk transcriptomic profiling through RNA-seq in quadriceps muscles from the same mice. Samples were collected at 4-hours after last drug/vehicle injection. Unbiased motif analysis showed the GR-binding element (GRE) motif as the top enriched motif in all groups **(fig2)** and peak tracks showed clear strong GR peaks upstream of the canonical GR marker *Fkbp5* **(fig2)**, indicating reliable GR ChIP-seq datasets for further quantitative comparisons.

**Figure 2.**
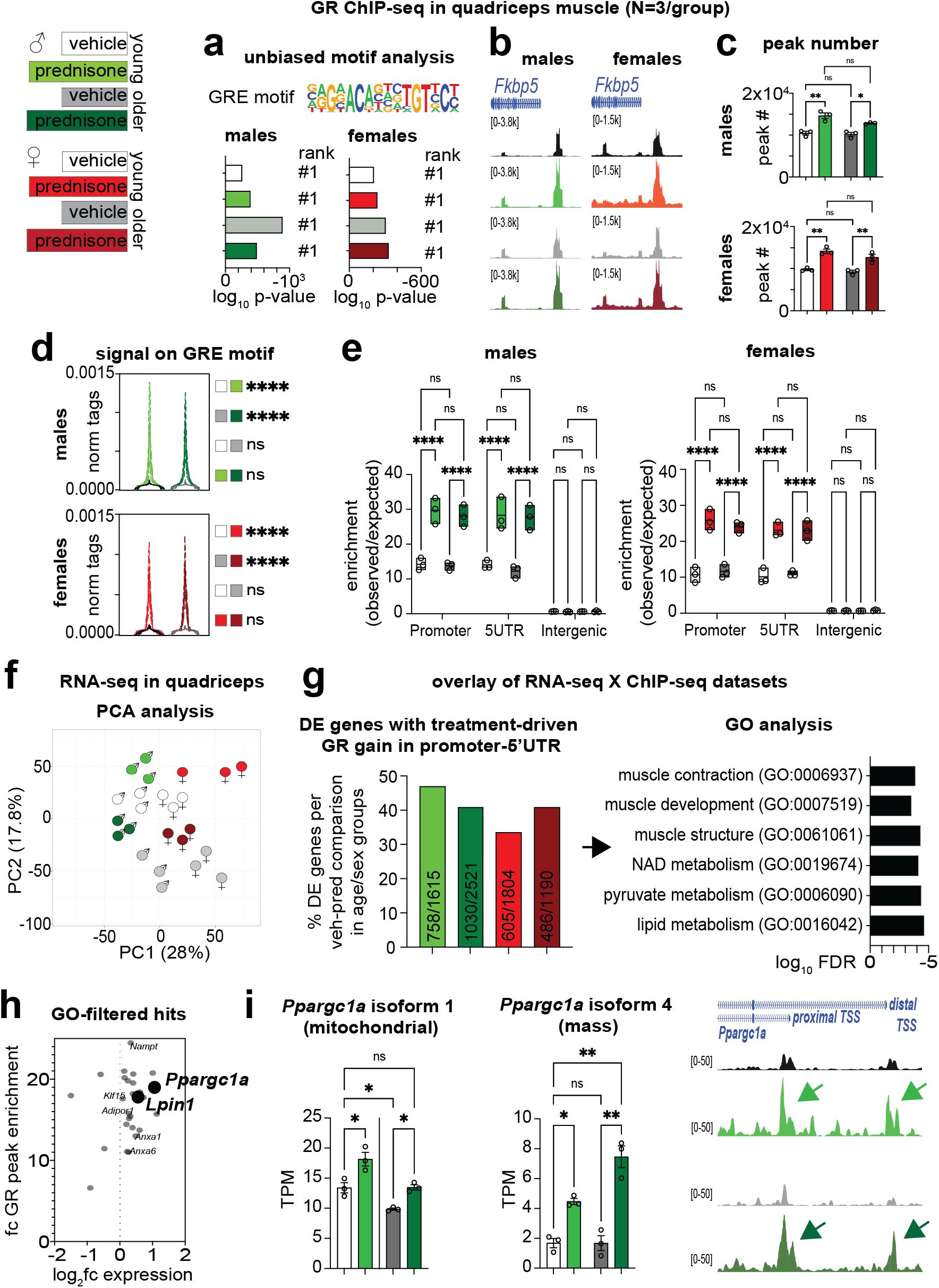
Epigenetic and transcriptional profiling reveals a treatment-induced muscle GR cistrome that is maintained through aging. **(a-b)** Motif analysis and robust promoter peaks in the canonical target *Fkbp5* confirm GR ChIP-seq datasets. **(c-e)** Treatment increased GR peak number and genome-wide, GRE-bound GR signal to comparable extents in both age (young, older) groups in both males and females. In all experimental groups, treatment increased GR signal in promoters and 5’UTR regions. **(f)** PCA analysis of RNA-seq datasets showed age- and treatment-related trends across sexes. **(g-h)** GO analysis revealed enrichment for muscle metabolic factors, particularly *Ppargc1a* (encoding PGC1alpha) and *Lpin1* (encoding the PGC1alpha co-factor Lipin1). **(i)** Expression of both isoforms 1 and 4 of *Ppargc1a* was rescued to young-like levels in treated older muscle, correlating with increased GR binding on canonical and alternative start sites (arrows). N=3/group; 2w ANOVA + Sidak: *, P<0.05; **, P<0.01; ***, P<0.001; ****, P<0.0001.

We first asked whether aging changed the muscle GR epigenomic activity in terms of peak number, signal on GREs and locus distribution at baseline and after treatment. Treatment increased GR peak number and average GR signal enrichment on the GRE motif genome-wide compared to vehicle in both young and aging muscle, but we did not find an age-specific effect in either vehicle- or treated-muscles **(fig2)**. Similarly, compared to vehicle, treatment increased GR signal in promoter-5’UTR rather than intergenic regions, but once again we did not find an age-specific effect in this shift **(fig2)**. Also, the trends were comparable in both male and female muscles **(fig2)**.

We then sought to overlay the GR ChIP-seq datasets with RNA-seq datasets to identify which GR targets were changing expression levels by treatment in both age groups in both sexes. Principal component analysis of the RNA-seq datasets showed overall sample clustering according to age, treatment and sex **(fig2)**. We over-layed GR ChIP-seq and RNA-seq following this question: how many/which genes are changed by treatment across age/sex groups with regards to differential RNA expression and increased GR signal in their promoter-5’UTR region?

We found that approximately 40% of the differentially expressed (DE) genes across all age/sex groups had increased GR signal in their promoter-5’UTR region. When compiling these gene lists together and analyzing them for pathway enrichment through gene ontology (GO), we found several pathways enriched related to muscle regulation and metabolism **(fig2)**. Considering the potential relevance of these pathways to the treatment-induced phenotype across aging, we used these GO pathways, i.e. the genes found enriched by GO analysis in these pathways, to filter out potential hits in play here. We confirmed several targets that we reported transactivated by intermittent prednisone in previous reports, including *Anxa1/6* ^31^, *Klf15* ^32^, *Nampt* ^29^ and *AdipoR1* ^26^ **(Suppl. Fig. 3A**).

However, we focused on the emergence of *Ppargc1a* (encoding PGC1alpha) and *Lpin1* (encoding Lipin1) among the top hits **(fig2)** based on two additional findings and literature-informed considerations. On one hand, running the isoform specific analyses from our paired-end RNA-seq datasets, we found that both the canonical mitochondria-regulating isoform 1 and the mass-regulating isoform 4 ^9^ were increased by treatment in both age groups and, consistent with the idea of double isoform transactivation, treatment increased GR peaks on both proximal (isoform 1) and distal (isoform 4) *Ppargc1a* TSS regions ^8^ in both young and aging muscle **(fig2; Suppl. Fig. 2B**). On the other hand, Lipin1 is a PGC1alpha co-factor and potentiates PGC1alpha activity through direct protein-protein interaction ^19^. We found that treatment increased GR transactivation of *Lpin1* in both ages, rescued the aging effect on *Lpin1* expression decrease in muscle **(Suppl. Fig. 3C**) and rescued the levels of PGC1alpha binding to Lipin1 **(Suppl. Fig. 3D**).

Thus, epigenomic and transcriptomic profiling identified a GR program that is elicited by intermittent prednisone and regulates muscle function and metabolism across aging. Intriguingly, we found a significant GR transactivation of PGC1alpha-Lipin1 and, in the next experiments, we sought to determine their role in the aging muscle rescue enabled by treatment.

### Muscle PGC1alpha is required by intermittent prednisone to coordinately stimulate energy and mass in muscle

To probe the extent to which PGC1alpha mediates the effects of chronic intermittent prednisone in muscle, we derived mice with myocyte-specific inducible deletion of PGC1alpha by crossing *Ppargc1a^fl/fl^*^33^ with *ACTA1-MerCreMer^+^* mice ^34^ on the C57BL/6J background. This background is slightly different than the one of the WT mice used in the young/aged cohorts, but i) it is consistent with all our other transgenic lines, including the Lipin1-KO used also in this study, and ii) the non-ablated control mice on this background recapitulated all the treatment features that we described above for mice on the 6JN background (see below). PGC1alpha ablation was induced starting at 3 months of age using intra-peritoneal (20 mg/kg per day for 5 days) and then chow-mediated intake (40mg/kg) of tamoxifen for 14 days, followed by 14 days of washout. These conditions allow us to reduce PGC1alpha levels in whole quadriceps muscle lysates by ∼85%, as we reported before ^29^. In this study, we compared *Cre^+/-^;Ppargc1a^wt/wt^* (PGC1alpha-WT) vs *Cre^+/-^;Pparcg1a^fl/fl^* (PGC1alpha-KO) male littermates after tamoxifen/washout to take into account both tamoxifen and Cre presence in both cohorts. After ablation/washout, mice were then started on 12-week-long regimens of intermittent prednisone/vehicle from 4 months of age, the same age/treatment conditions used in the young cohorts of the previous experiment. We used this timeline to minimize the adult muscle adaptations to gene ablation before treatment effects. Considering the epigenomic/transcriptomic screening of the initial cohorts took into account targets common to both sexes, in these subsequent mechanistic experiments we focused on only one sex (males) to maintain power to detect trends while decreasing the overall number of mice.

The *Pparcg1a*-floxed allele features *loxP* sites surrounding exons 3-5, which are shared by the transcripts of both isoforms 1 and 4 ^9^. We therefore verified that both transcripts were deleted in our PGC1alpha-KO muscles at baseline. Compared to PGC1alpha-WT, PGC1alpha-KO muscle showed profound downregulation of total *Ppargc1a*, as well as isoform1 and isoform4 transcripts **(fig3)**, as assessed through qPCR using previously reported discriminating primers ^9^. Regarding overall function, despite no differences induced by genotype at baseline and endpoint (i.e. WT vehicle vs KO vehicle), PGC1alpha ablation blocked the treatment effect in both grip strength and treadmill tests **(fig3)**. Regarding force production, PGC1alpha ablation did not change specific force while did decrease fatigue resistance in vehicle-treated mice, and further blocked the treatment effect on increases in both parameters **(fig3)**.

**Figure 3.**
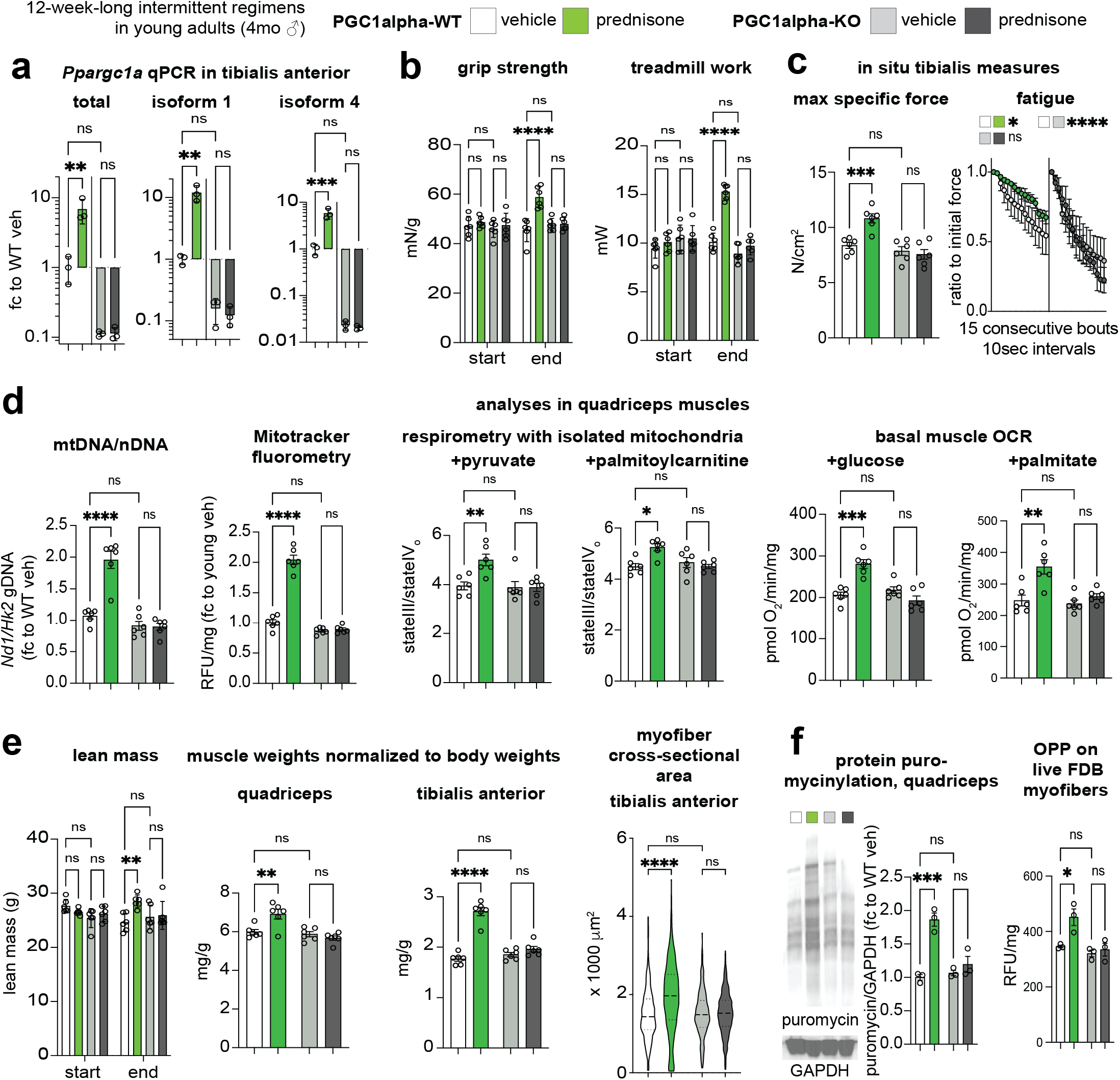
Myocyte-specific inducible PGC1alpha ablation blocks treatment effects on both mitochondrial function and muscle mass. **(a)** Recombination of the floxed allele reduced expression of both PGC1alpha isoforms in muscle. **(b-c)** In young adult mice, myocyte-specific inducible PGC1alpha ablation blocked the effects of 12-week-long intermittent prednisone treatment on strength, treadmill, force and fatigue. **(d)** PGC1alpha ablation blocked or blunted treatment effects on mitochondrial abundance and on mitochondrial RCR and basal OCR in muscle tissue regardless of fuel. **(e)** PGC1alpha ablation blocked or blunted treatment effects on lean mass, muscle mass, myofiber CSA. **(f)** Treatment increased protein translation in muscle dependent on myocyte PGC1alpha. N=3-6/group; 2w ANOVA + Sidak: *, P<0.05; **, P<0.01; ***, P<0.001; ****, P<0.0001.

Previously, we found that myocyte PGC1alpha is required for the mitochondrial effects of a single circadian-specific glucocorticoid pulse ^29^. We therefore checked whether this was still true for the chronic intermittent prednisone treatment effects on muscle mitochondria. Indeed, consistent with the finding of PGC1alpha transactivation by chronic treatment, PGC1alpha ablation blocked the treatment effect on mitochondrial abundance, respiratory control ratio of isolated mitochondria and basal oxygen consumption of quadriceps muscle tissue **(fig3)**. The KO effects on respiration appeared related to overall mitochondrial function rather than shifts in nutrient preference, as they were recapitulated with either glucose/pyruvate or palmitate/palmitoylcarnitine as fuels **(fig3)**.

Strikingly and unexpectedly, PGC1alpha ablation also blocked the treatment effects on lean and muscle mass. Despite no genotype differences in vehicle-treated mice, PGC1alpha ablation blocked the treatment effect on lean mass, muscle/body weight ratio (shown here for both muscles used for analyses in this model) and myofiber cross-sectional area **(fig3)**. Moreover, the treatment/KO effects appeared related to anabolic propensity as shown by two independent puromycin-based assays of protein synthesis: protein puromycinylation in quadriceps muscle lysates after in vivo puromycin injection, and ex vivo O-propargyl-puromycin incorporation/fluorometry ^35^ in live myofibers **(fig3)**.

Thus, inducible PGC1alpha ablation in adult myocytes without long-term and/or developmental adaptations blocks the effects of chronic intermittent prednisone not only on mitochondrial capacity, but also muscle mass.

### PGC1alpha mediates the treatment effect on increased carbon shuttling between oxidative intermediates and amino acids in muscle

We were intrigued by the fact that the myocyte-specific PGC1alpha mediated both mitochondrial and mass effects of treatment. We therefore hypothesized that the upregulated PGC1alpha mediated a myocyte-autonomous metabolic program coordinating the increased nutrient oxidation with anabolic growth. Amino acid availability determines mass rescue in sarcopenia ^36^. Intermediary metabolites like pyruvate and TCA cycle intermediates are direct precursors of amino acids like alanine, glutamine and aspartate, generally decreased in aging muscle ^37^. We therefore asked whether PGC1alpha coordinated the increased glucose oxidation in treated muscle with amino acid biogenesis. To investigate this in our transgenic muscles, we traced the glucose contribution to in-muscle amino acids using an ex vivo system where the isolated muscle undergoes repetitive contractions in the presence of ^13^C-labeled glucose and insulin ^32^. Albeit not in steady-state, this system offers the advantage of quantitating muscle-autonomous effects without circulating and extra-muscle contributions, and we previously used it to trace macronutrient fate, including glucose, after intermittent prednisone treatments in dystrophic muscles ^32^ and normal versus obese WT muscles ^26^.

We compared PGC1alpha-WT/-KO muscles after vehicle/prednisone treatment for labeling rates of oxidative intermediates (pyruvate, alpha-ketoglutarate, oxaloacetate) and their putative amino acid products (alanine, glutamate/glutamine, aspartate) downstream of uniformly labeled ^13^C_6_-glucose. We also quantitated labeling rates for amino acids produced from glycolytic intermediates (serine, glycine). For each metabolite, we quantitated overall fractional labeling i.e. percentage of the sum of all ^13^C-labeled isoforms versus the total sum of labeled and unlabeled isoforms. Compared to vehicle controls, treated PGC1alpha-WT muscles showed increased labeling of the oxidative intermediates and their amino acid products, but the treatment effect was blocked or blunted with myocyte PGC1alpha ablation **(fig4)**. Also, no significant treatment or genotype effects were quantitated for labeled serine and glycine **(fig4)**, and PGC1alpha ablation did not significantly impact the treatment-driven increase in overall glucose tolerance **(Suppl. Fig. 4A**), further underscoring the mitochondrial specificity of this regulation rather than a change in overall glucose utilization.

**Figure 4.**
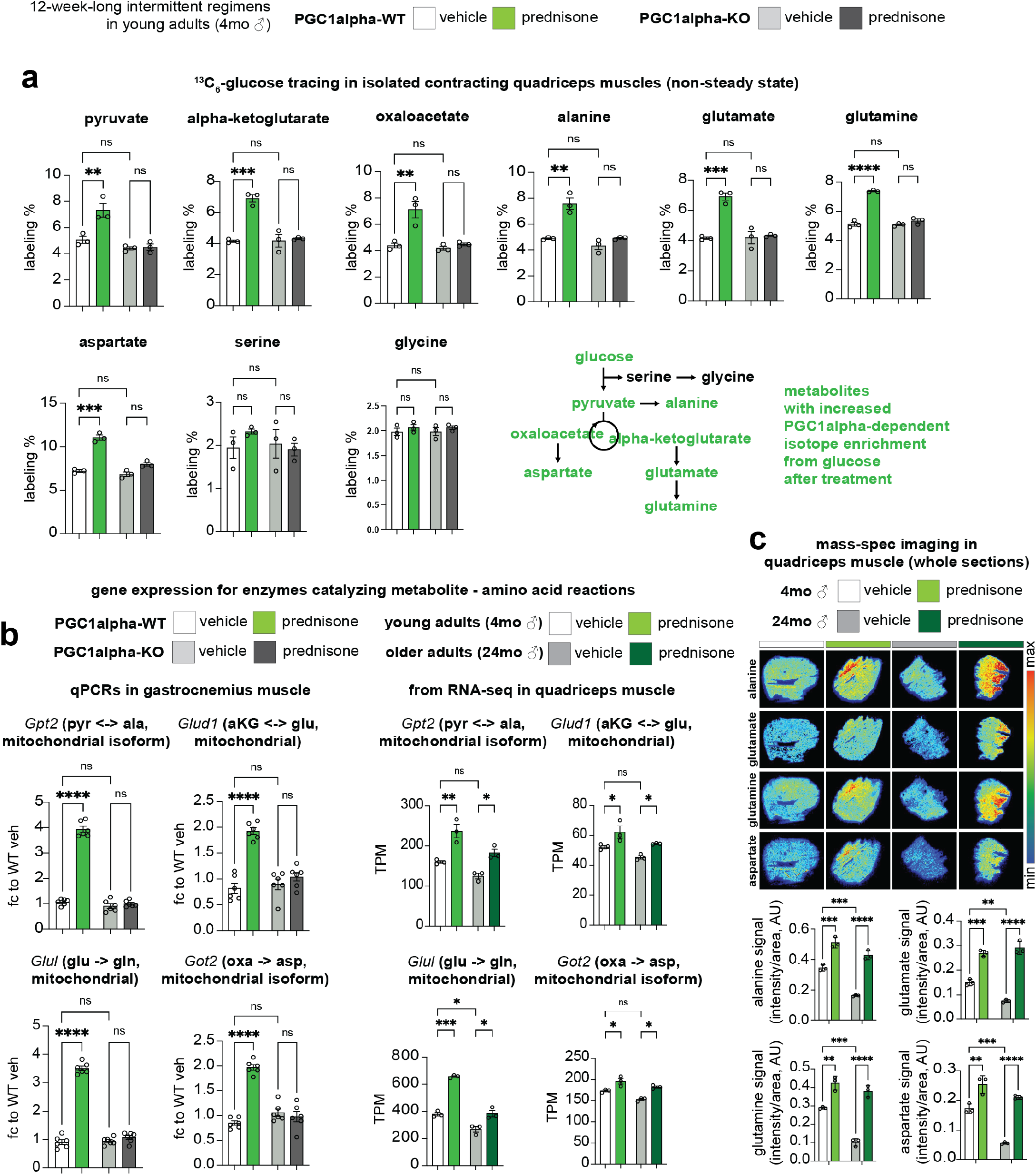
Treatment increases carbon shuttling between glucose oxidation and amino acids in muscle dependent on myocyte-specific PGC1alpha. **(a)** In isolated contracting muscle exposed to ^13^C_6_-glucose, treatment increased overall labeling ratio of amino acids through oxidative intermediates (alanine, glutamate, glutamine, aspartate), but not through glycolytic intermediates (serine, glycine). **(b)** Myocyte-specific PGC1alpha was required for treatment-driven upregulation of mitochondrial enzymes and/or enzyme isoforms mediating the underlying reactions between glucose metabolites and amino acids. The treatment effect on expression of those genes was also confirmed in young and older muscles per RNA-seq. (c) Mass-spec imaging showed increased levels of target amino acids in treated young and aged muscles after a glucose+insulin challenge. N=3-6/group; 2w ANOVA + Sidak: *, P<0.05; **, P<0.01; ***, P<0.001; ****, P<0.0001.

In light of a prior report implicating PGC1alpha in the transcriptional control of the mitochondrial alanine transaminase^38^ (gene name, *Gpt2*), we asked whether we could quantitate a PGC1alpha-dependent effect on the mitochondrial enzymes or mitochondrial enzyme isoforms mediating the carbon shuttling between the metabolites and amino acids found enriched in labeling. Therefore, in addition to *Gpt2* (pyruvate <-= alanine), we quantitated expression levels of *Glud1* (alpha-ketoglutarate <-> glutamate), *Glul* (glutamate -> glutamine), *Got2* (oxaloacetate -> aspartate) in the gastrocnemius muscles of the same animals whose quadriceps muscles were used for the glucose tracing experiments. For all those four enzymes, PGC1alpha ablation blocked or blunted the treatment-driven upregulation seen in PGC1alpha-WT muscle **(fig4, left; Suppl. Fig. 4B**). Accordingly, we checked against our RNA-seq datasets in young/older mice and found that the same four enzyme genes were upregulated by treatment in muscles of both young and older age groups **(fig4, right)**. We tested whether the treatment effect on glucose-derived amino acid biogenesis was quantifiable in young and aged muscle. At 24 hours after last treatment injection, we challenged young and aged mice with 1g/kg glucose coupled with 0.5U/kg insulin to maximize muscle glucose uptake across ages and treatment groups. Mass-spec imaging on cryosections from quadriceps muscles collected at 2-hours post-challenge showed increased levels of alanine, glutamate, glutamine and aspartate after treatment in both young and aged muscles **(fig4)**.

Thus, the myocyte-specific PGC1alpha mediates the metabolic program enabled by intermittent prednisone in muscle to coordinate increased nutrient oxidation with amino acid biogenesis.

### Myocyte-specific Lipin1 is required for the pro-ergogenic effects of treatment upstream of PGC1alpha

Considering the combined upregulation of *Ppargc1a* isoforms 1 and 4 by treatment, we asked whether each isoform was sufficient to rescue a specific parameter of treatment effect, i.e. mitochondrial abundance by isoform 1 and muscle mass by isoform 4. We generated AAVs to overexpress either GFP (control), or *Ppargc1a* isoform 1, or isoform 4 downstream of a *CMV* promoter. Adult myocyte tropism was promoted by using the MyoAAV serotype^39^ and confirmed via qPCR in WT muscle tissue at 2 weeks after a single r.o. injection of 10^12^vg/mouse (**fig5, left)**. We then used PGC1alpha-KO mice for a genetic rescue experiment with AAV-overexpression of isoform 1, isoform 4 or both, in vehicle versus treatment conditions. On one hand, isoform 1 was sufficient to enable the treatment effect on mitochondrial abundance but not mass, consistent with the impact of this isoform on oxidative efficiency^40^. On the other hand, isoform 4 was sufficient to enable the treatment effect on muscle mass but not mitochondrial abundance, consistent with prior reports^9^. Importantly, treatment showed a significant additive effect over the genetic rescue effect **(fig5, right)**. In addition to our inducible KO data, this rescue experiment confirmed the specific roles of PGC1alpha isoforms 1 and 4 in the combined energy-mass effect of intermittent prednisone. However, the additive effect of treatment over AAV re-expression made us hypothesize that an additional factor induced by treatment is required for fully coaxing the PGC1alpha upregulation to the global anti-sarcopenic effect.

**Figure 5.**
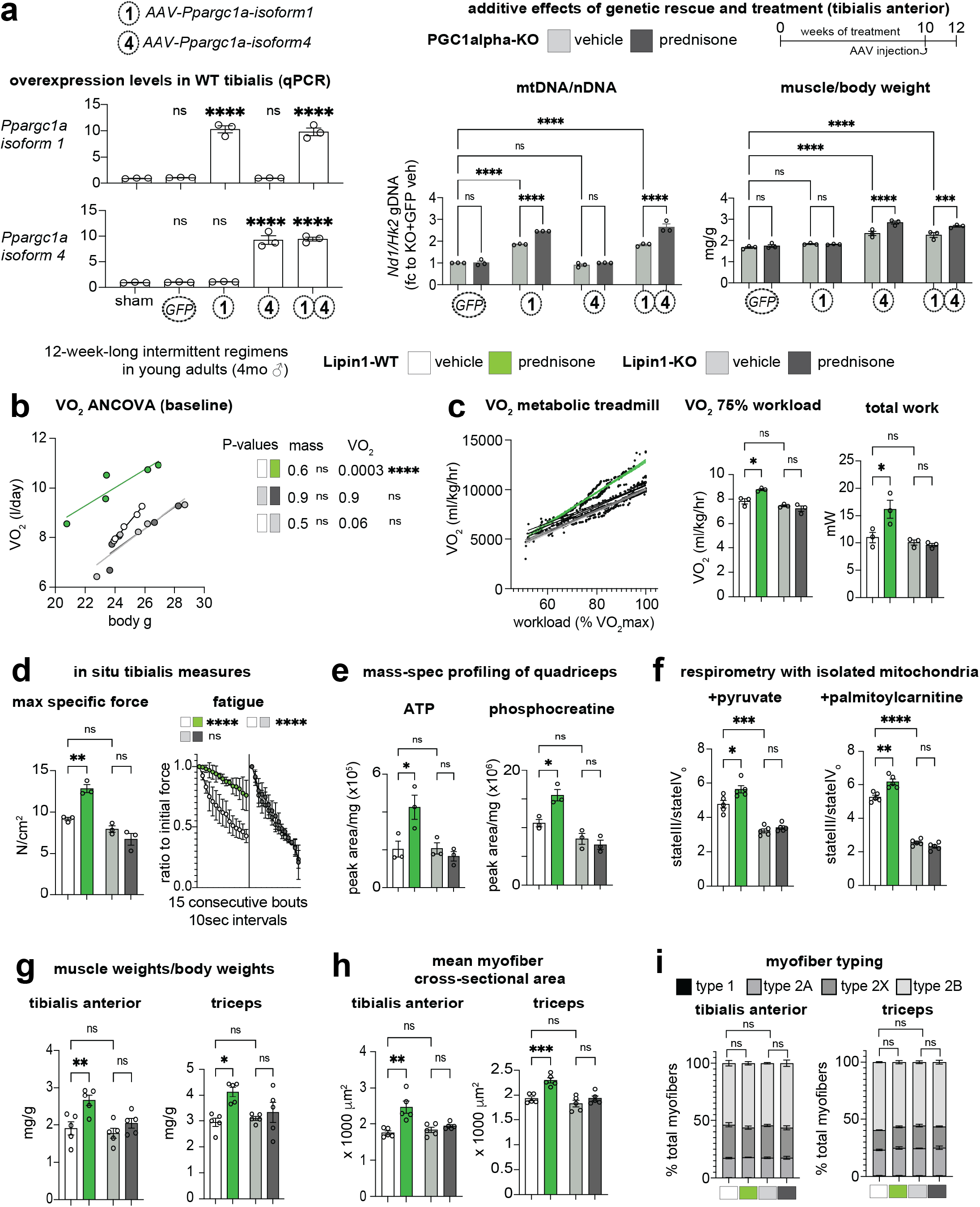
Myocyte-specific Lipin1 controls energy-mass balance in muscle. **(a)** MyoAAV-mediated overex-pression in WT muscle at 2 weeks post r.o. injection of 10^12vg/mouse (left). Combination of AAV and treatment in PGC1alpha-KO mice revealed an additive effect of treatment on genetic rescues of mitochondrial abundance by Pgc1alpha isoform 1 and muscle mass by Pgc1alpha isoform 4 (right). Together with our RNA-seq/ChIP-seq screening, the additive effect warranted investigation of Lipin1 as treatment-driven co-factor to coax PGC1alpha regulation with energy-mass balance. **(b)** ANCOVA analysis for VO2 in metabolic cages without specific exercise triggers showed increased VO_2_ independent from body mass in control mice (Lipin1-WT), but not after Lipin1 ablation (Lipin1-KO). **(c)** In the metabolic treadmill, treatment increased VO2 at 75% workload and overall work at exhaustion dependent on myocyte-specific Lipin1. **(d-f)** Lipin1 was critical for treatment-driven effects on muscle force and fatiguability, ATP and phosphocreatine, and mitochondrial respiration. **(g-i)** Analogously to its co-factor PGC1alpha manipulation, Lipin1 ablation blunted or blocked treatment effects on muscle mass in two different locomotory muscles (tibialis, hindlimbs; triceps, forelimbs) without sizeable changes in myofiber typing. N=3-5/group; 2w ANOVA + Sidak: *, P<0.05; **, P<0.01; ***, P<0.001; ****, P<0.0001.

Within that hypothesis, we were intrigued by the emergence of Lipin1 as GR transactivation target with intermittent prednisone in tandem with PGC1alpha in aging muscle. Indeed, prior experiments in liver showed that Lipin1 is a direct co-activator of PGC1alpha and primes it to enhance its pro-metabolic gene program ^19^. Constitutive knockout of Lipin1 impairs muscle function and mitochondrial metabolism in young and mature adult ages ^16^. In the context of intermittent prednisone effects, we reasoned that, if PGC1alpha mediates energy-mass co-ordination and Lipin1 is a critical PGC1alpha activator, Lipin1 could also be required for treatment effects.

We generated transgenic mice for myocyte-restricted inducible ablation of Lipin1 by crossing *Lpin1^fl/fl^* (floxed exon7, complete protein loss ^16^) with *ACTA1-MerCreMer^+^*mice ^34^ on the C57BL/6J background. We used the same tamoxifen/washout protocol as PGC1alpha-KO and found ∼85% Lipin1 ablation in quadriceps muscle **(Suppl. Fig. 5A**). Consistent with the PGC1alpha-KO experiments, we compared *Cre^+/-^;Lpin1^wt/wt^* (Lipin1-WT) vs *Cre^+/-^;Lpin1^fl/fl^*(Lipin1-KO) male littermates for the effects of a 12-week-long regimen of intermittent prednisone/vehicle from 4 months of age.

Considering the potential involvement of Lipin1 in muscle oxidative capacity, we interrogated the extent to which the myocyte-specific Lipin1 regulated body-wide VO_2_ at rest and during aerobic exercise in metabolic cages and treadmill. Indeed, aging-related exercise intolerance is evident in mice through a decrease in the slope of the VO_2_/workload curve and in the VO_2_ rates at baseline, maximal (100%) and also submaximal (75%) workloads ^41^. At rest, intermittent prednisone increased VO_2_ independent from body mass, while inducible Lipin1 ablation induced a downward trend in the vehicle-treated muscles and further blocked the treatment effect **(fig5)**. Analogously, during acute exercise on the treadmill, treatment significantly increased the VO_2_/workload curve slope, the average VO_2_ at 75% workload and the overall treadmill work in Lipin1-WT but not Lipin1-KO mice **(fig5)**.

In parallel to VO_2_ and exercise capacity trends, Lipin1 ablation blocked the treatment effect on specific force and fatigue, decreasing the resistance to fatigue also in vehicle-treated muscle **(fig5)**. Also, Lipin1 deletion blocked the treatment effect on ATP and phosphocreatine levels, as well as on mitochondrial respiratory control ratio **(fig5)**. Intriguingly and in line with our PGC1alpha findings and Lipin1-PGC1alpha protein-protein interaction findings, Lipin1 ablation also blocked the treatment effect on the muscle mass parameters of muscle/body weight ratios and myofiber cross-sectional area in two different locomotory muscles (tibialis, hind limbs; triceps, fore limbs) without sizable shifts in myofiber typing **(fig5)**. These effects were obtained even though Lipin1 ablation did not significantly change the treatment effect on muscle PGC1alpha upregulation compared to vehicle control **(Suppl. Fig. 5B**).

Thus, Lipin1 is a GR transactivation target in aging muscle by intermittent prednisone and its inducible postnatal ablation recapitulates the PGC1alpha-mediated role on energy-mass coordination, underscoring Lipin1 as key factor in the treatment-PGC1alpha muscle program.

## Discussion

In aggregate, our data show that exogenous glucocorticoid intermittence can rescue both mitochondrial and mass defects in the aging muscle through the PGC1alpha-Lipin1 axis. Our study unveils several points of novelty along with intriguing open questions regarding glucocorticoid signaling and muscle remodeling in aging.

A remarkable point brought forward by these experiments is the fact that the aging muscle maintained sensitivity to the pro-ergogenic impact of prednisone intermittence. Here we are using the term “pro-ergogenic” to refer to improved exercise tolerance accompanied by increases in both mitochondrial function and muscle mass. Despite analogous trends in dystrophic ^32^ and obese muscle ^26^, the question of whether aging blunts or skews the muscle responsiveness to intermittent prednisone remained open. Indeed, aging is known to strongly undermine the intrinsic susceptibility of muscle to both anabolic and mitochondrial stimuli ^42,43^. In that regard, it was remarkable to identify several molecular markers of an age- and sex-independent (i.e. maintained from young to older muscles in both males and females) program enabled by treatment, including extent of epigenomic GR activation and non-canonical targets of GR transactivation like PGC1alpha and Lipin1. Based on the functional improvements recapitulated by treatment to a variable extent in aging mice of both sexes, we focused here on “inclusive” gene programs underlying comparable remodeling across ages and sexes. However, because sex-specific differences can be identified in specific molecular markers of intermittent prednisone effects in young adult mice ^44^, future studies are warranted to better investigate how aging affects the “exclusive”, sex-dimorphic programs enabled by glucocorticoid intake.

Our findings linking the pro-ergogenic GR program to simultaneous upregulation of both “mitochondrial” and “mass” *Pparcg1a* isoforms argues in favor of two additional concepts to the puzzling role of PGC1alpha in muscle aging. On one hand, our data demonstrate that a balanced upregulation of both isoforms promotes the balanced rescue of both mitochondrial capacity and muscle mass in the context of sarcopenia. On the other hand, the GR engagement on both isoform TSSs by the glucocorticoid regimen we used here (once-weekly 1mg/kg prednisone at ZT0) implicates the myocyte GR as context-specific “additional factor” that coaxes the PGC1alpha isoform regulation with muscle remodeling outputs. Regarding both aspects, our data here pave the way to several compelling questions for the aging muscle, including shared/differential mechanisms of PGC1alpha isoforms and which co-factors are engaged by the GR in beneficial versus deleterious muscle contexts. It is interesting to emphasize that our findings are consistent with not only genetic experiments with PGC1alpha isoforms^8–10,40^ but also with scattered reports regarding the metabolic link between PGC1alpha activation and growth. For instance, PGC1alpha was found to integrate oxidative and growth pathways downstream of caloric restriction ^11^, a generally positive intervention in the context of aging and muscle health. Constitutive transgenic PGC1alpha overexpression in muscle changed not only TCA cycle but also amino acid levels ^45^, suggesting a role for this factor in coordinating mitochondrial intermediates with amino acids. A prior example in that regard comes from a study in C2C12 myoblasts that implicated PGC1alpha in increasing pyruvate-alanine interconversion through transcriptional control over *Gpt2*, the mitochondrial alanine aminotransferase ^38^. The role of PGC1alpha in mitochondrial proteostasis has emerged as an important determinant of muscle health and exercise efficacy^6^. Future studies are warranted to pinpoint in significant granularity the effects of prednisone intermittence on the many key aspects of proteostasis in sarcopenia^46^.

Furthermore, our study identifies an additional non-redundant factor required for coaxing PGC1alpha activation towards energy-mass rescue in sarcopenia, i.e. Lipin1. Our data implicate *Lpin1* as GR transactivation target in muscle, a regulation that so far was only reported in adipose tissue ^47^. Moreover, our transgenic mice data involve Lipin1 as the required co-factor to link PGC1alpha to the pro-ergogenic program enabled by glucocorticoid intermittence in the aging muscle. This provides an intriguing new context for the understudied Lipin1 property as PGC1alpha co-activator, which was investigated until now only in liver ^19^. Considering the recent findings with Lipin1 ablation and complex lipid metabolism in heart ^48^, future studies are warranted in better identifying how Lipin1 regulates lipotoxicity in the sarcopenic muscle, particularly in light of the recent complex role reported for Lipin1 in counteracting dystrophic pathophysiology ^20^.

Glucose metabolism contributes to cell mass during development ^49^ and – as highlighted by our data – also in aging. The role of the mitochondrial TCA cycle in providing the “building blocks’ for many anabolic pathways is well known ^50^, yet the advantages and shortcomings of this metabolic hub in the aging muscle still need further elucidation. For instance, a likely important question to address in the future will be how the glucocorticoid intermittence regulates the relationship between non-essential and essential amino acids availability, an important point for mass regulation and exercise tolerance in the aging muscle ^51^. An intriguing new angle in that perspective is laid out by the chrono-pharmacology underpinnings of the prednisone injection time we used here, ZT0 (lights-on). The early light-phase in mice is marked by the daily trough in the main rodent endogenous glucocorticoid, corticosterone ^52^, and the daily peak of epigenetic activity by the activating clock factors BMAL1/CLOCK ^53^. Aging alters endogenous glucocorticoid rhythmicity in both mice and humans ^54^, and the fact that the pro-functional GR program was consistently elicited in both young and older mice in our experiments shifts the focus of attention from corticosterone/cortisol rhythm to the myocyte-autonomous circadian system. Indeed, the muscle-intrinsic circadian rhythm is a fundamental regulator of large swaths of muscle homeostasis and changes with aging ^55^. We already reported that ZT0 is a critical circadian time-of-intake to coax the muscle GR to transiently interact with the clock factor BMAL1 and promote pro-functional, non-canonical transcriptional programs ^29^. However, the extent to which aging undermines or maintains the time-specific glucocorticoid effects on the muscle clock is still unknown and deserves further studying, particularly with regards to consequential outputs like insulin sensitivity, amino acid availability, bioenergetic management. This will be key to progress discovery and translation of our initial findings here for the treatment of sarcopenia. A supportive piece of in-patient evidence in that sense is a recent pilot clinical trial in a heterogenous cohort of dystrophic patients that reported positive effects of intermittent prednisone with 7-9pm intake time (approximately analogous to ZT0 in mice) on both lean mass and muscle function ^27^. Nonetheless, further studies are required to not only define the clock requirements of such effects with aging, but also to identify the evidence-based biomarkers to reliably elicit such effects in the aging human organism.

### Limitations of the study

Our study presents important limitations to keep in mind when interpreting our findings, especially when projecting their translational potential. We used here aging WT mice between 24 and 27 months of age as main model of sarcopenia. While recapitulating several key hallmarks of aging ^56^, aging mice do not fully mimic the extraordinarily complex biology of human aging. Also, an important consideration for our mechanistic studies with PGC1alpha and Lipin1 is that our experiments with myocyte-specific inducible ablation were conducted at 4 months of age (young adulthood). This is based on the fact that our most promising functional treatment phenotypes and epigenomic-transcriptomic screening hits in the aging cohorts were recapitulated in the young adult cohorts too. However, we recognize that this is most likely an oversimplification and that age-matched ablation studies, i.e. ablation at 24 months of age, will shed light over additional molecular mechanisms in play here.

In summary, our study reports on the thought-provoking phenotypes and muscle-autonomous mechanisms of anti-sarcopenic action by exogenous glucocorticoid intermittence. Our findings challenge the current paradigm on glucocorticoids and muscle regulation, opening a potential new perspective on the signals and the muscle effectors that are actionable to counteract aging-related exercise intolerance and strength loss.

## Acknowledgements

NMR profiling was performed thanks to the Cincinnati Children’s NMR Metabolomics Facility (RRID: SCR_022636), with critical assistance by Drs. Romick and Watanabe. Mass-spec analyses were performed thanks to the Mass-Spec Metabolomics Core Facility at Robert H. Lurie Comprehensive Cancer Center of Northwestern University, with critical assistance by Dr. Gao. Next-gen sequencing was performed thanks to the Cincinnati Children’s DNA Sequencing and Genotyping Facility (RRID: SCR_022630), with critical assistance by David Fletcher, Keely Icardi, Julia Flynn and Taliesin Lenhart.

## Grant support

This work was supported by R01AG078174-01, R01HL166356-01, R03DK130908-01A1 (NIH) and RIP, CCRF Endowed Scholarship, HI Translational Funds (CCHMC) grants to MQ; R01HL119225-01A1 (NIH) to BF.

## Materials and Methods

### Animal handling and treatments

Mice were housed in a pathogen-free facility in accordance with the American Veterinary Medical Association (AVMA) and under protocols fully approved by the Institutional Animal Care and Use Committee (IACUC) at Cincinnati Children’s Hospital Medical Center (#2022-0020, #2023-0002). Consistent with the ethical approvals, all efforts were made to minimize suffering. Euthanasia was performed through carbon dioxide inhalation followed by cervical dislocation and heart removal. Mice were maintained on a 14h/10h light/dark cycle, approximately 22°C constant temperature, and ad libitum access to chow and water. Aged (24 months-old at treatment start) and young control (4 months-old at treatment start) male and female cohorts were obtained from the NIA Aged Rodent Colony, C57BL/6JN background. Mice for mechanistic studies were obtained and interbred from Jackson Laboratories (Bar Harbor, ME) and/or colleagues. PGC1alpha-KO mice and -WT littermates from crossing #025750 and #009666 lines, Cre^-^ and Cre^+^ littermates obtained from *Ppargc1a^fl/wt^ x Ppargc1a^wt/fl^;HSA-MerCreMer^+/-^* matings. Analogously, we obtained Lipin1-KO mice and -WT littermates from crossing *Lpin1^fl/fl^* (floxed exon7, complete protein loss ^16^) with the HSA-MCM line. Both PGC1alpha- and Lipin1-KO mice were on the C57BL/6J background. Gene ablation was induced starting at 3 months of age using intra-peritoneal (20 mg/kg per day for 5 days; Sigma #T5648) and then chow-mediated intake (40 mg/kg; Harlan #TD.130860) of tamoxifen for 14 days, followed by 14 days of washout ^29^. Weekly prednisone treatment consisted of once-weekly i.p. injection of 1mg/kg prednisone (#P6254; Sigma-Aldrich; St. Louis, MO) ^32^. The injectable solution was diluted from a 5mg/ml stock in DMSO (#D2650; Sigma-Aldrich; St. Louis, MO) in 50 ml volume. Injections were conducted at the beginning of the light-phase (ZT0; lights-on). For epigenetic analyses, mice were sacrificed, and tissues harvested at 4-hours (ZT4) after last prednisone injection in chronic or single-pulse treatments. For non-epigenetic-involving experiments, tissues were harvested 24 hours after last prednisone injection in chronic or single-pulse treatments, i.e., ZT0. All in vivo, ex vivo and post-mortem analyses were conducted blinded to treatment groups.

### Analyses of muscle function, lean and muscle mass, myofiber typing

Our routine procedures concerning body composition, muscle function, mass and myofiber typing can be found as point-by-point protocols here ^57^.

Forelimb grip strength was monitored using a meter (#1027SM; Columbus Instruments, Columbus, OH) blinded to treatment groups. Animals performed ten pulls with 5 seconds rest on a flat surface between pulls. Grip strength was expressed as force normalized to body weight. Running endurance was tested on a motorized treadmill with electrified resting posts (#1050RM, Columbus Instruments, Columbus, OH) and 10° inclination. Speed was accelerated at 1m/min^2^ starting at 1m/min and individual test was interrupted when the subject spent >30sec on resting post. Running endurance was analyzed as weight-normalized cumulative work (mW)^58^. Immediately prior to sacrifice, in situ tetanic force from tibialis anterior muscle was measured using a Whole Mouse Test System (Cat #1300A; Aurora Scientific, Aurora, ON, Canada) with a 1N dual-action lever arm force transducer (300C-LR, Aurora Scientific, Aurora, ON, Canada) in anesthetized animals (0.8 l/min of 1.5% isoflurane in 100% O2). Specifications of tetanic isometric contraction: initial delay, 0.1 sec; frequency, 200Hz; pulse width, 0.5 msec; duration, 0.5 sec; stimulation, 100mA ^32^. Muscle length was adjusted to a fixed baseline of ∼50mN resting tension for all muscles/conditions. Force-frequency cruve was measured from 25 Hz to 200 Hz with intervals of 25 Hz, pause 1 minute between tetani. Fatigue analysis was conducted by repeating tetanic contractions every 10 seconds until complete exhaustion of the muscle (50 cycles). Specific force was calculated (N/mm^2^) for each tetanus frequency as (P_0_ N)/[(muscle mass mg/1.06 mg/mm^3^)/L_f_ mm]. 1.06 mg/mm^3^ is the mammalian muscle density. L_f_=L_0_*0.6, where 0.6 is the muscle to fiber length ratio in *tibialis anterior* muscle ^59^. We reported here specific force values in N/cm^2^ units.

Myofiber cross-sectional areas were obtained from histology analyses at endpoint. Excised tissues were fixed in 10% formaldehyde (Cat #245-684; Fisher Scientific, Waltham, MA) at room temperature for ∼24 hours, then stored at ^+^4C before processing. Seven mm sections from the center of paraffin-embedded muscles were stained with hematoxylin and eosin (H&E; cat #12013B, 1070C; Newcomer Supply, Middleton, WI). CSA quantitation was conducted on >400 myofibers per tissue per mouse. Imaging was performed using an Axio Observer A1 microscope (Zeiss, Oberkochen, Germany), using 10X and 20X (short-range) objectives. Images were acquired through Gryphax software (version 1.0.6.598; Jenoptik, Jena, Germany) and quantitated through ImageJ ^60^.

Magnetic resonance imaging (MRI) scans to determine lean mass ratios (% of total body mass) were conducted in non-anesthetized, non-fasted mice at ZT8 using the EchoMRI-100H Whole Body Composition analyzer (EchoMRI, Houston, TX). Mice were weighed immediately prior to MRI scan. Before each measurement session, system was calibrated using the standard internal calibrator tube (canola oil). Mice were scanned in sample tubes dedicated to mice comprised between 20 g and 40 g body mass. Data were collected through built-in software EchoMRI version 140320. Data were analyzed when hydration ratio > 85%.

Muscle mass was calculated as muscle weight immediately after sacrifice and explant, normalized to whole body weight. For myofiber typing, sections were incubated with primary antibodies BA-F8 (1:10), SC-71 (1:30) and BF-F3 (1:10; all by Developmental Studies Hybridoma Bank, Iowa City, IA) overnight at 4°C. Then, sections were incubated with secondary antibodies AlexaFluor350 anti-IgG2b, AlexaFluor488 anti-IgG1 and AlexaFluor594 anti-IgM (Cat #A21140, A21121, 1010111; Life Technologies, Grand Island, NY). Type 1 fibers stained blue, type 2A stained green, type 2X showed no staining, type 2B stained red. Myofiber types were then quantitated over at least five serial sections and quantitated as % of total counted myofibers.

All analyses were conducted blinded to treatment.

### Respirometry with isolated mitochondria and muscle tissue

Basal tissue OCR values were obtained from basal rates of oxygen consumption of muscle biopsies at the Seahorse XF HS Mini Extracellular Flux Analyzer platform (Agilent, Santa Clara, CA) using previously detailed conditions ^32^. Basal OCR was calculated as baseline value (average of 3 consecutive reads) minus value after rotenone/antimycin addition (average of 3 consecutive reads). Basal OCR values were normalized to total protein content, assayed in each well after the Seahorse run through homogenization and Bradford assay. Nutrients: 5 mM glucose, 1 mM palmitate-BSA (#G7021, #P0500; Millipore-Sigma, St Louis, MO); inhibitors: 0.5 mM rotenone ^+^ 0.5 mM antimycin A (Agilent).

Respiratory control ratio (RCR) values were obtained from isolated mitochondria from muscle tissue. Quadriceps are harvested from the mouse and cut up into very fine pieces. The minced tissue is placed in a 15mL conical tube (USA Scientific #188261) and 5 mL of MS-EGTA buffer with 1 mg Trypsin (Sigma #T1426-50MG) is added to the tube. The tube is quickly vortexed and the tissue is left submerged in the solution. After 2 minutes, 5 mL of MS-EGTA buffer with 0.2% BSA (Goldbio #A-421-250) is added to the tube to stop the trypsin reaction. MS-EGTA buffer: Mannitol-ChemProducts #M0214-45, Sucrose-Millipore #100892, HEPES-Gibco #15630-080, EGTA-RPI #E14100-50.0. The tube is inverted several times to mix then set to rest. Once the tissue has mostly settled to the bottom of the tube, 3 mL of buffer is aspirated and the remaining solution and tissue is transferred to a 10mL glass tissue homogenizer (Avantor # 89026-382). Once sufficiently homogenized the solution is transferred back into the 15mL conical tube and spun in the centrifuge at 1,000 × g for 5 minutes at 4 degrees Celsius. After spinning, the supernatant is transferred to a new 15 mL conical tube. The supernatant in the new tube is then centrifuged at 12,000 × g for 10 minutes at 4 degrees Celsius to pellet the mitochondria. The supernatant is discarded from the pellet and the pellet is then resuspended in 7 mL of MS-EGTA buffer and centrifuged again at 12,000 × g for 10 minutes at 4 degrees Celsius. After spinning, the supernatant is discarded, and the mitochondria are resuspended in 1 mL of Seahorse medium (Agilent #103335-100) with supplemented 10 µL of 5 mM pyruvate (Sigma #P2256-100G) and 10 µL of 5 mM malate (Cayman Chemical #20765). After protein quantitation using a Bradford assay (Bio-Rad #5000001), 2.5 µg mitochondria are dispensed per well in 180 µl total volumes and let to equilibrate for 1 hour at 37°C. 20 µL of 5 mM ADP (Sigma #01905), 50 µM Oligomycin (Milipore #495455-10MG), 100 µM Carbonyl cyanide-p-trifluoromethoxyphenylhydrazone (TCI #C3463), and 5 µM Rotenone (Milipore #557368-1GM)/Antimycin A (Sigma #A674-50MG) are added to drug ports A, B, C, and D respectively to yield final concentrations of 500, 50, 10, and 0.5 µM. Nutrients: 0.5mM pyruvate, 0.1 mM palmitoylcarnitine (#P2256, #61251; Millipore-Sigma, St Louis, MO). At baseline and after each drug injection, samples are read for three consecutive times. RCR was calculated as the ratio between state III (OCR after ADP addition) and uncoupled state IV (OCR after oligomycin addition). Seahorse measurements were conducted blinded to treatment groups.

### Mitochondrial density, NMR and mass-spec profiling

The mtDNA/nDNA assay was performed on genomic DNA isolated using the Gbiosciences Omniprep kit (Gbiosciences #786-136). The ratio was obtained from qPCR values (absolute expression normalized to internal standard Rn45s and DNA concentration) of ND1 (mtDNA locus; primers: CTAGCAGAAACAAAC-CGGGC, CCGGCTGCGTATTCTACGTT) versus HK2 (nDNA locus; primers: GCCAGCCTCTCCTGATTTTAG-TGT, GGGAACACAAAAGACCTCTTCTGG)^61^. For the Mitotracker assay, Mitotracker Green FM powder (Invitrogen #M7514) is resuspended in 373 µL of DMSO (Fisher #BP231-100) to obtain a 200 µM concentration. One microliter of this resuspension is added to 1 mL of Mammalian Ringer’s Solution (Electron Microscopy Sciences #11763-10) containing isolated myofibers from the flexor digitorum brevis muscle (FDB) of the mouse foot. The solution containing myofibers and Mitotracker is then pipetted into a 96 well plate (Corning #9017) in increments of 200 µL. This plate is then read at the plate reader for fluorescence with excitation set to 490nm and emission set to 516 nm. Values are then normalized to protein content, assayed in each well after the Mitotracker assay through homogenization and Bradford assay.

NMR profiling was performed at the NMR Metabolomics Facility at CCHMC. Each individual sample of quadriceps muscle tissue was homogenized using a cryogenic ball-mill homogenizer (Cryomill, Retsch Inc.) and weighted into bead tubes (2.8 mm, MO BIO, catalog #13114-50) for extraction. The average water content of skeletal muscle tissue was determined prior to this experiment which was 78%. Based on the weights of tissues, the solvent volumes used for extraction were calculated for each sample. Modified Bligh and Dyer extraction^62–64^ was used to obtain polar metabolites. Briefly, cold methanol and water were added to the samples in bead tubes and homogenized for 30 s. The samples were transferred into glass tubes containing cold chloroform and water. The final methanol:chloroform:water ratio is 2:2:1.8. The mixture was vortexed, incubated on ice for 10 min, then centrifuged at 2000x *g_n_* for 5 min. The polar phase was transferred into 1.5mL Eppendorf tube and dried by vacuum centrifuge for 4-5 hrs at room temperature. The dried metabolites were resuspended in 220 µL of NMR buffer containing 100 mM phosphate buffer, pH7.3, 1 mM TMSP (3-Trimethylsilyl 2,2,3,3-d4 propinoate), 1mg/mL sodium azide) prepared in D_2_O. Final volume of 200 µL of each sample was transferred into 3 mm NMR tube (Bruker) for data collection. One-dimensional ^1^H NMR spectra were acquired on a Bruker Avance III HD 600 MHz spectrometer with 5 mm, BBO Prodigy probe. All data were collected at a calibrated temperature of 298 K using the noesygppr1d pulse sequence in the Bruker pulse sequence library. All the data collection and processing were performed using Topspin 3.6 software (Bruker Analytik, Rheinstetten, Germany). For a representative sample, two-dimensional data ^1^H-^1^H total correlation spectroscopy (TOCSY) and 2D ^1^H-^13^C heteronuclear single quantum coherence (HSQC) were collected for metabolites assignment. Chemical shifts were assigned to metabolites based on 1D ^1^H, 2D TOCSY and HSQC NMR experiments with reference spectra found in databases, Human Metabolome Database (HMDB)^65^, and Chenomx® NMR Suite profiling software (Chenomx Inc. version 8.1). The concentrations of the metabolites in polar extracts were calculated using Chenomx software based on the internal standard, TMSP (1 mM). The metabolites concentrations were normalized to the original tissue weights for further analysis.

Mass-spec profiling of hydrophilic metabolites was performed at the Metabolomics Mass-spec Core of Northwestern University. Total hydrophilic metabolite content was extracted from quadriceps muscle tissue at treatment endpoint through methanol:water (80:20) extraction, adapting conditions described previously ^66^. Briefly, total metabolite content from quadriceps muscle was obtained from ∼100 mg (wet weight) quadriceps muscle tissue per animal. Frozen (-80°C) muscle was pulverized in liquid nitrogen and homogenized with ∼250 µl 2.3 mm zirconia/silica beads (Cat # 11079125z, BioSpec, Bartlesville, OK) in 1 ml methanol/water 80:20 (vol/vol) by means of Mini-BeadBeater-16 (Cat # 607, Biospec, Bartlesville, OK) for 1 minute. After centrifuging at 5000 rpm for 5 minutes, 200 µl of supernatant were transferred into a tube pre-added with 800 µL of icecold methanol/water 80% (vol/vol). Samples were vortexed for 1 min, and then centrifuged at ∼20,160 × *g* for 15 min at 4°C. Metabolite-containing extraction solution was then dried using SpeedVac (medium power). 200 µl of 50% Acetonitrile were added to the tube for reconstitution following by overtaxing for 1 min. Samples solution were then centrifuged for 15 min @ 20,000 × g, 4°C. Supernatant was collected for LC-MS analysis for Hydrophilic Metabolites Profiling as follows. Samples were analyzed by High-Performance Liquid Chromatography and High-Resolution Mass Spectrometry and Tandem Mass Spectrometry (HPLC-MS/MS). Specifically, system consisted of a Thermo Q-Exactive in line with an electrospray source and an Ultimate3000 (Thermo) series HPLC consisting of a binary pump, degasser, and auto-sampler outfitted with a Xbridge Amide column (Waters; dimensions of 4.6 mm × 100 mm and a 3.5 µm particle size). The mobile phase A contained 95% (vol/vol) water, 5% (vol/vol) acetonitrile, 20 mM ammonium hydroxide, 20 mM ammonium acetate, pH = 9.0; B was 100% Acetonitrile. The gradient was as following: 0 min, 15% A; 2.5 min, 30% A; 7 min, 43% A; 16 min, 62% A; 16.1-18 min, 75% A; 18-25 min, 15% A with a flow rate of 400 μL/min. The capillary of the ESI source was set to 275 °C, with sheath gas at 45 arbitrary units, auxiliary gas at 5 arbitrary units and the spray voltage at 4.0 kV. In positive/negative polarity switching mode, an m/z scan range from 70 to 850 was chosen and MS1 data was collected at a resolution of 70,000. The automatic gain control (AGC) target was set at 1 × 10^6^ and the maximum injection time was 200 ms. The top 5 precursor ions were subsequently fragmented, in a data-dependent manner, using the higher energy collisional dissociation (HCD) cell set to 30% normalized collision energy in MS2 at a resolution power of 17,500. The sample volumes of 25 µl were injected. Data acquisition and analysis were carried out by Xcalibur 4.0 software and Tracefinder 2.1 software, respectively (both from Thermo Fisher Scientific). Metabolite levels were analyzed as peak area normalized to wet tissue weight (weight before cryo-pulverization). Metabolite analysis was performed blinded to treatment groups.

### Isotope tracing

^13^C tracing from nutrients in muscle was performed adapting reported conditions^67^ to our muscle stimulus settings used to probe muscle force (see below). Immediately after sacrifice, quadriceps muscles were dissected and immobilized on a Sylgard-coated well of a 12-multiwell plate by means of two 27g needles at the muscle extremities. The well was pre-filled with 1X Ringers’ solution (146 mM NaCl, 5 mM KCl, 2 mM CaCl2, 1 mM MgCl2, 10 mM HEPES; pH, 7.4) containing 25 mU/ml insulin (Cat #RP-10908; Thermo Fisher, Waltham, MA) and 10 mM U-^13^C_6_-glucose (Sigma #310808) and kept at 37°C on a heated pad. The nutrient solution was constantly bubbled with a 95%O_2_/5%CO_2_ line (∼2psi). After 5 min equilibration in solution, electrodes were inserted at the muscle extremities, securing them to the holder needles. Using a Whole Mouse Test System (Cat #1300A; Aurora Scientific, Aurora, ON, Canada), 20 contractions (1X/min) were induced with following specifications: initial delay, 0.1 sec; frequency, 200Hz; pulse width, 0.5 msec; duration, 0.5 sec; 100 mA stimulation. Muscles were then removed from the isotope-containing solution, quickly rinsed in nutrient-free Ringers’ solution, dried and immediately flash-frozen. Muscle metabolites were then extracted and analyzed as per metabolomics procedures (LC-MS, see above), and mass resolution was carried on pre-determined metabolites, while control energetics (ATP, phosphocreatine) were analyzed from simultaneous quantitation from the LC-MS system. Metabolite labeling ratio was calculated on peak area/mg tissue values subtracting the background ^13^C labeling ratio obtained from muscles exposed to unlabeled nutrients (same reagents used for respirometry), and expressed as labeling % per metabolite, i.e. (sum labeled isoforms)/(sum labeled+unlabeled isoforms). Metabolite analysis was performed blinded to treatment groups.

### Mass-spec imaging, i.e. matrix-assisted laser desorption/ionization (MALDI) mass spectrometry

Frozen quadriceps muscles were cryosectioned into 12 μm-thick cryosections using a Leica CM1860 cryostat. Cryosections were mounted onto indium tin oxide coated glass slides (Delta Technologies Limited, Loveland, Colorado), then coated with 2′,4′,6′-Trihydroxyacetophenone monohydrate at 10 mg/ml in 50:50 cyclohexane/methanol using a HTX TM Sprayer. 20 passes were performed over each tissue at a spray volume of 50 ml/min and nozzle temperature of 50°C. Once sprayed, samples were individually wrapped in plastic bags and immediately transferred to -80°C. MALDI MS imaging was performed using a Q-Exactive HF mass spectrometer (Thermo Scientific, Bremen, DE) fitted with a MALDI/ESI Injector (Spectroglyph LLC, Kennewick, WA). Laser post-ionization (MALDI-2) was used to enhance analytical sensitivity for triglycerides and cholesteryl esters. Images were acquired at 20 micrometer voxel size, using a pulse energy of ∼6 mJ and repetition rate of 30Hz. Q Exactive HF MS Scan parameters were optimized for polar metabolites: polarity – negative, scan range – 350-1500 m/z, resolution – 120,000, automatic gain control – off, maximum inject time 250 ms. ImageInsight™ (Spectroglyph LLC) software was used for initial data visualization and to convert data files into imzML format for visualization and further processing in SciLS™ software (Bruker, Billerica, MA). All metabolite images produced were normalized to the total ion chromatogram.

### Metabolic cages and metabolic treadmill

VO_2_ in baseline conditions (ml/h; expressed as aggregate values of l/day) was assessed via indirect calorimetry using the Prometheon Automated Phenotyping System (Sable Systems International, Las Vegas, NV) at the shared Metabolic Cage facility in the CCHMC Vet Services. Data collection started at 24 hours after last prednisone or vehicle injection and lasted for 5 days. Results are expressed as average values (all mice per group, all values per mouse, average of 5 days) over a circadian period, as well as in an ANCOVA analysis (test for difference in regression lines; performed through CalR^68^) with average values of active phase plotted against body mass values per mouse, as recommended by ^69^.

For VO_2_ analysis during aerobic exercise, we used an Oxymax Metabolic Treadmill (Columbus Instruments, Columbus, OH), using the stepwise speed increase protocol described previously to separate young vs aged mice based on the slope of the VO_2_/workload curve and VO_2_ rates at baseline, submaximal and maximal (75%) workloads ^41^. Treadmill belt was angled 10° uphill to match our regular treadmill conditions and calculate work. Mice were assessed at the metabolic treadmill at 24hours after last vehicle or prednisone injection.

Metabolic cage and metabolic treadmill assessments were performed blinded to regimens or genotype.

### ChIP-sequencing and RNA-sequencing

Quadriceps muscles were cryopowdered using a liquid nitrogen-cooled Retsch Cryomill. The cryopowdered tissue was then fixed in 10 ml 1% PFA for 30 minutes at room temperature with gentle nutation. Fixation was quenched 1ml of 1.375M glycine (Cat # BP381-5, Fisher Scientific, Waltham, MA) with gentle nutation for 5 minutes at room temperature. After centrifugation at 3000g for 5 minutes at 4°C, the pellet was resuspended in cell lysis buffer as per reported conditions ^70^, supplementing the cell lysis buffer with 3mg/ml cytochalasin B and rotating for 10 minutes at 4°C. Nuclei were pelleted at 300g for 10 min at 4°C, and subsequently processed following the reported protocol with the adjustment of adding 3µg/ml cytochalasin B into all solutions for chromatin preparation and sonication, antibody incubation, and wash steps. Chromatin was then sonicated for 15 cycles (30 sec, high power; 30 sec pause; 500µl volume) in a water bath sonicator set at 4°C (Bioruptor 300; Diagenode, Denville, NJ). After centrifuging at 10000g for 10 minutes at 4°C, sheared chromatin was checked on agarose gel for a shear band comprised between ∼150 and ∼600bp. Two µg of chromatin was kept for pooled input controls, whereas ∼50µg chromatin were used for each pull-down reaction in a final volume of 2ml rotating at 4°C overnight. Primary antibody: rabbit polyclonal anti-GR (Abclonal #A2164). Chromatin complexes were precipitated with 100µl Dynabeads M-280 (sheep anti-rabbit #11203D; Thermo Scientific, Waltham, MA). After washes and elution, samples were treated with proteinase K (cat #19131; Qiagen, Hilden, Germany) at 55°C and cross-linking was reversed through overnight incubation at 65°C. DNA was purified using the MinElute purification kit (cat #28004; Qiagen, Hilden, Germany), quantitated using Qubit reader and reagents. Library preparation and sequencing were conducted at the CCHMC DNA sequencing Core. Up to 30 ng of post-Immunoprecipitation dsDNA as measured by Invitrogen™ Qubit™ highsensitivity spectrofluorometric measurement was made into Illumina-compatible NGS libraries using the ThruPLEX® DNA-Seq kit from Takara Bio Inc. Each created library was dual-indexed with 8-base molecular barcodes for high level mul5plexing. After 13 cycles of PCR amplification, completed libraries were sequenced on an Illumina NovaSeq™ model (6000 or X Plus) sequencer, generating 20 million or more high quality, 100 base length read pairs per sample. A quality control check on the generated fastq files was performed using FastQC. Peak analysis was conducted using HOMER software (v4.10, ^71^; standard commands) after aligning fastq files to the mm10 mouse genome using bowtie2 ^72^. PCA was conducted using ClustVis ^73^. Heatmaps of peak density were imaged with TreeView3 ^74^. Peak tracks were imaged through WashU epigenome browser. Gene ontology pathway enrichment was conducted using the Gene Onthology analysis tool ^75^.

RNA-seq was conducted on RNA extracted from quadriceps muscle. Total RNA was extracted from cryo-pulverized quadriceps muscles with Trizol (#15596026; Thermo Fisher Scientific, Waltham, MA) and re-purified using the Rneasy Mini Kit (Cat #74104; Qiagen, Germantown, MD). RNA-seq was performed at the DNA Core (CCHMC) blinded to treatment groups. 150 to 300 ng of total RNA determined by Qubit (Cat#Q33238; Invitrogen, Waltham, MA) high-sensitivity spectrofluorometric measurement was poly-A selected and reverse transcribed using Illumina’s TruSeq stranded mRNA library preparation kit (Cat# 20020595; Illumina, San Diego, CA). Each sample was fitted with one of 96 adapters containing a different 8 base molecular barcode for high level multiplexing. After 15 cycles of PCR amplification, completed libraries were sequenced on an Illumina NovaSeqTM 6000, generating 20 million or more high quality 100 base long paired end reads per sample. A quality control check on the fastq files was performed using FastQC. Upon passing basic quality metrics, the reads were trimmed to remove adapters and low-quality reads using default parameters in Trimmomatic ^76^ [Version 0.33]. The trimmed reads were then mapped to mm10 reference genome using default parameters with strandness (R for single-end and RF for paired-end) option in Hisat2 ^77^ [Version 2.0.5]. In the next step, transcript/gene abundance was determined using kallisto ^78^ [Version 0.43.1]. We first created a transcriptome index in kallisto using Ensembl cDNA sequences for the reference genome. This index was then used to quantify transcript abundance in raw counts and counts per million (CPM). Differential expression (DE genes, FDR<0.05) was quantitated through DESeq2 ^79^. PCA was conducted using ClustVis ^73^. Heatmaps were imaged with TreeView3 ^74^. Gene ontology pathway enrichment was conducted using the Gene Onthology analysis tool ^75^.

### WB, qPCR, OPP assays

Protein analysis was performed on ∼50 mg total lysates from whole quadriceps muscles homogenized in general protein buffer, i.e., PBS supplemented with 1 mM CaCl_2_, 1 mM MgCl_2_ (#C1016, #M8266, Sigma-Aldrich; St. Louis, MO) and protease and phosphatase inhibitors (#04693232001, #04906837001, Roche, Basel, Switzerland). Blocking and stripping solutions: StartingBlock and RestorePLUS buffers (#37543, #46430, ThermoFisher Scientific, Waltham, MA). Co-immunoprecipitation was performed from ∼100 mg protein lysates, which were rotated overnight at 4°C in a final volume of 500 µl protein buffer with 5 µl primary antibody for pull-down. The day after, 50µl Dynabeads were added to the samples, with additional rotating incubation at 4°C for 4 hours. After 4 washes at the magnet separator, proteins were extracted from Dynabeads through incubation at 95°C for 15 minutes in Laemmli buffer. Input controls were run as 10 mg of the input protein lysates. Primary antibodies (all diluted 1:1000 for O/N incubation at ^+^4°C): rabbit anti-PGC1alpha (ABClonal #A12348), rabbit anti-GR (ABClonal #A2164), mouse anti-puromycin (cat#PMY-2A4; DSHB, Iowa City, IA), rabbit anti-GLUD1 (ABClonal #A7631), rabbit anti-GLUL (ABClonal #A21856), rabbit anti-GOT2 (ABClonal #A19245), rabbit anti-GPT2 (ABClonal #A23670), rabbit anti-LIPIN1 (ABClonal #A14111). Secondary antibodies (diluted 1:5000 for 1-hour incubation at room temperature): HRP-conjugated donkey anti-rabbit or anti-mouse (#sc-2313 and #sc-2314, Santa Cruz Biotech, Dallas, TX). Counterstain for loading control was performed with ponceau (#P7170, Sigma-Aldrich; St. Louis, MO). Blots were developed with SuperSignal Pico (cat#34579; Thermo Scientific, Waltham, MA) using the iBrightCL1000 developer system (cat #A32749; Thermo Scientific, Waltham, MA) with automatic exposure settings. WB gels and membranes were run/transferred in parallel and/or stripped for multiple antibody-based staining for densitometry analyses. Protein density was analyzed using the Gel Analysis tool in ImageJ software ^60^ and expressed as fold changes to control samples.

OPP fluorometry was performed adapting the regular instructions for the Click-iT™ Plus OPP Alexa Fluor™ 488 Protein Synthesis Assay Kit (cat #C10456; Thermo Scientific, Waltham, MA) to live myofibers, isolated from the flexor digitorum brevis (FDB) muscle using previously reported conditions^80^. Protein puromycinylation was assessed in gastrocnemius muscle tissue through anti-puromycin WB of whole protein lysates at 30 min post-i.p. puromycin injection (0.040 μmol/body g; #P8833 Sigma-Aldrich; St. Louis, MO).

For RT-qPCR assays, total RNA was extracted from cryo-pulverized quadriceps muscles with Trizol (#15596026; Thermo Fisher Scientific, Waltham, MA) and 1 µg RNA was reverse-transcribed using 1X qScript Supermix (#95048; QuantaBio, Beverly, MA). RT-qPCRs were conducted in triplicates using 1X Sybr Green Fast qPCR mix (#RK21200, ABclonal, Woburn, MA) and 100nM primers at a CFX96 qPCR machine (Bio-Rad, Hercules, CA; thermal profile: 95°C, 15 sec; 60°C, 30 sec; 40X; melting curve). Primers were selected among validated primer sets from the MGH Primer Bank; IDs: GPT2-27805389a1; GLUD1-6680027a1; GLUL-31982332a1; GOT2-6754036a1; LIPIN1-27923941a1; and from published primers sets: PGC1a1 and PGc1a4 primers ^9^; mitochondrial DNA quantification primers ^61^.

### AAV preparation and injection

Approximately 70-80% confluent HEK293T cells (AAVpro® 293T Cell Line; Takara # 632273 AAVpro® 293T Cell Line; Takara # 632273) in DMEM (SH30022.01, Cytiva Life Sciences) supplemented with 2% Bovine Growth Serum (BGS; Cytiva Life Sciences), and 1.0 mM Sodium Pyruvate were triple transfected with pHelper (Cell Biolabs; #340202), pAAV-GOI (Vector Builder; (VB230317-1361ncv; pAAV[Exp]-CMV>mPpargc1a[NM_008904.3]*-V5:WPRE); (VB230317-1364xmj; pAAV[Exp]-CMV>mPpargc1a_isoform4-Myc:WPRE)) and pAAV Rep-Cap (1A-Myo; re-cloned from published sequence^39^, gift from Molkentin lab) plasmids using PEI, Linear, MW250,000 (PolySciences, Inc) in 40-T150mm cell culture plates. Eighteen hours after transfection, medium is changed to DMEM supplemented with 1% BGS, 1.0 mM Sodium Pyruvate, and 1X MEM Non-essential Amino Acid Solution (Sigma; M7148). Approximately 96 hours post-transfection, the media and cells were collected and processed separately. Cells were lysed using repeated freeze/thaw cycles at a minimum of five times in 1X Gradient Buffer (0.1 M Tris, 0.5 M NaCl, 0.1 M MgCl_2_). The cell debris were then treated with Benzonase Endonuclease at 0.65 μL per 5 mL (Sigma-Aldrich #1037731010 (100000 Units)) for at least one hour. The homogenates were cleared from debris by centrifugation. AAVs were precipitated from the cell medium with polyethylene glycol (PEG) 8000 The PEG-precipitated AAV was collected by centrifugation, and the AAV pellet was resuspended in 1X GB. Media and cell AAV’s were combined and loaded onto an Iodixanol (OptiPrep Density Gradient Medium; Sigma-Aldrich #D1556250) gradient at 15%, 25%, 40% and 60% in 1X Gradient Buffer, and subject to ultracentrifugation. The 40% iodixanol layer, containing the AAV particles, was extracted and a buffer exchange into 2xPBS/10mM MgCl2 was performed using Centrifugal Filters (30000 NMWL (30K), 4.0 mL Sample Volume; Millipore-Sigma #UFC803024, and 100000 NMWL (100K), 15.0 mL Sample Volume; Millipore-Sigma # UFC910024). Primers binding within the AAV-GOI ITR’s CMV region (Forward: GTTCCGCGTTACATAACTTACGG; Reverse: CTGCCAAGTGGGCAGTTTACC) were used to measure the virus titer with quantitative polymerase chain reaction (qPCR). Before releasing the viral DNA from the particles, all extra-viral DNA was removed by digestion with DNase I. Then, the viral DNA was released by Proteinase K digestion. For injection, 10^12vg was diluted in 50 μl saline and injected r.o. in anesthetized mice (isoflurane 1.5%) the same day of prednisone injection at week 10 of treatment.

### Statistics

Statistical analyses were performed using Prism software v9.2.0 (Graphpad, La Jolla, CA). The Pearson-D’Agostino normality test was used to assess data distribution. When comparing data groups for more than one related variable, two-way ANOVA was used with Sidak multi-comparison (treatment vs age effect; treatment vs KO effect). Significance scores reported on charts: *, P<0.05; **, P<0.01; ***, P<0.001; ****, P<0.0001. When data points < 10, data were presented as single values (dot plots, histograms). Tukey distribution bars or violin plots were used to emphasize data range distribution for > 10 data points per pool. For curves, the s.e.m. values for each plotted point were reported as upper and lower lines.

---

**Supplementary Figure 1.**
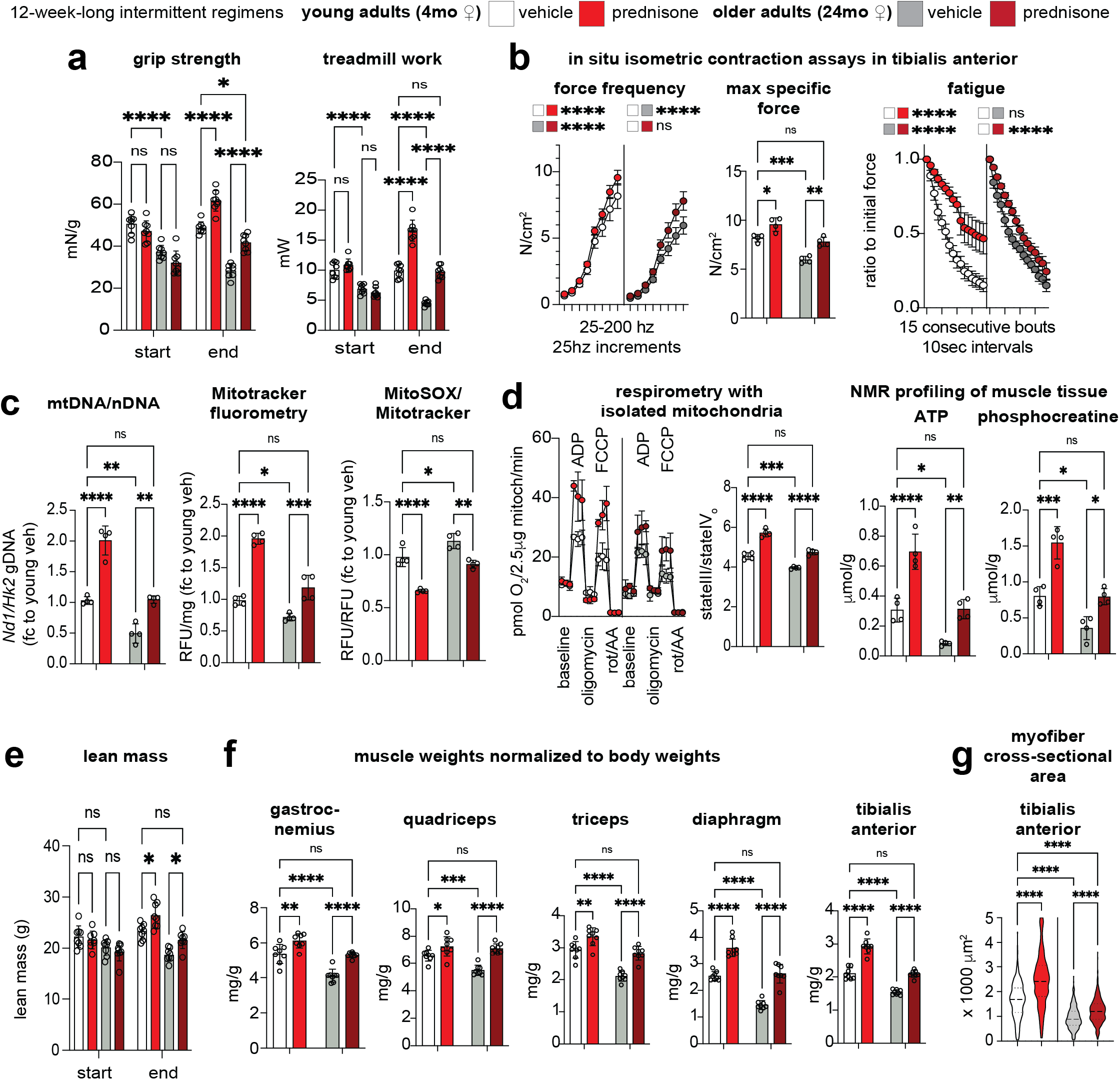
Related to Figure 1. Additional analyses in young/older female mice. Parallel analyses in age- and background-matched WT females showed analogous treatment effects on overall function and muscle energy-mass, as compared to the trends in male mice shown in Figure 1. N=4-8/group; 2w ANOVA + Sidak: *, P<0.05; **, P<0.01; ***, P<0.001; ****, P<0.0001.

**Supplementary Figure 2.**
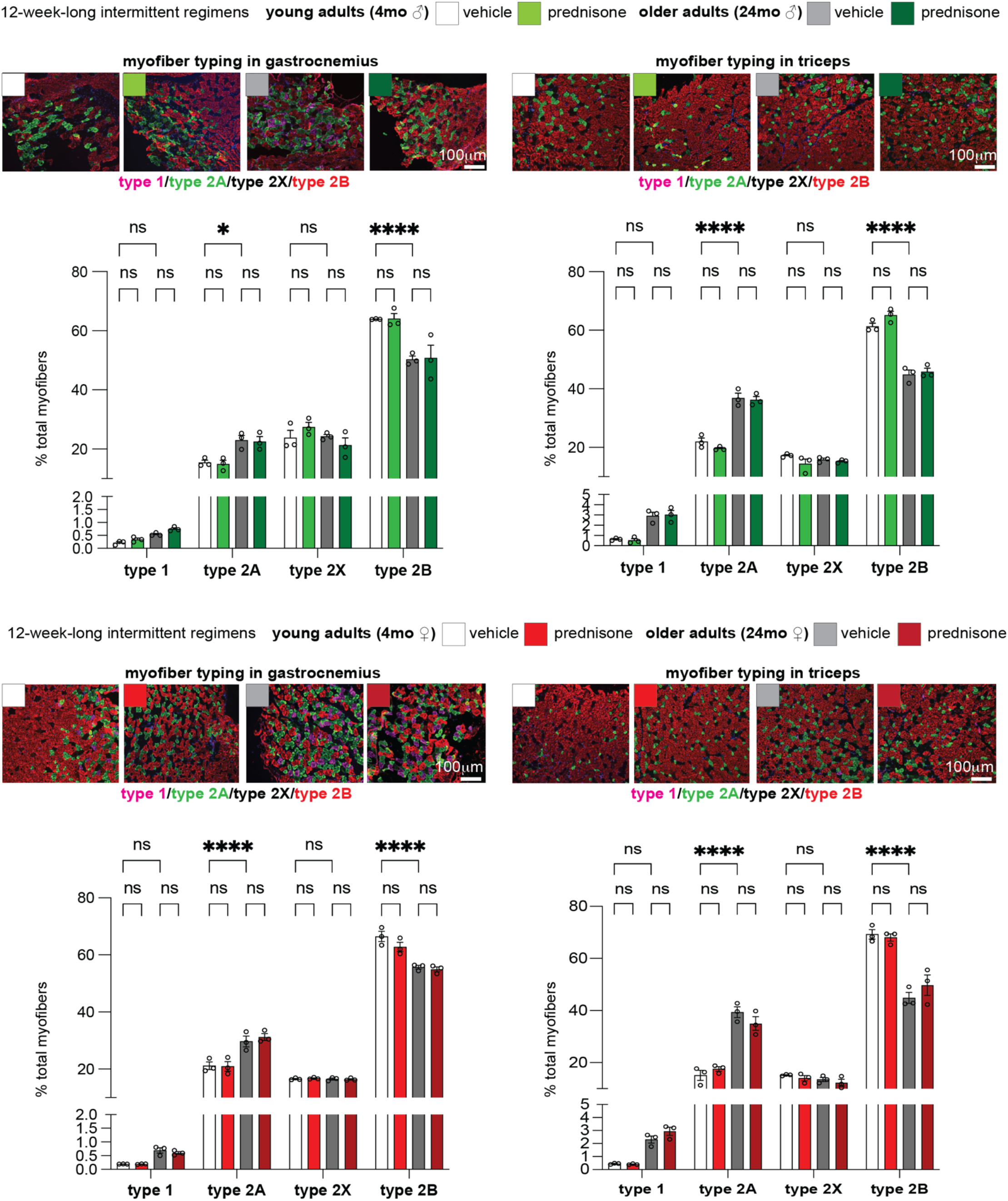
Related to Figure 1. Treatment did not sizably change myofiber type distribution. No significant effects of treatment were observed on top of the expected age-related shifts in both male and female mice in two locomotory muscles, gastrocnemius (hindlimbs) and triceps (forelimbs). N=3/group; 2w ANOVA + Sidak: *, P<0.05; **, P<0.01; ***, P<0.001; ****, P<0.0001.

**Supplementary Figure 3.**
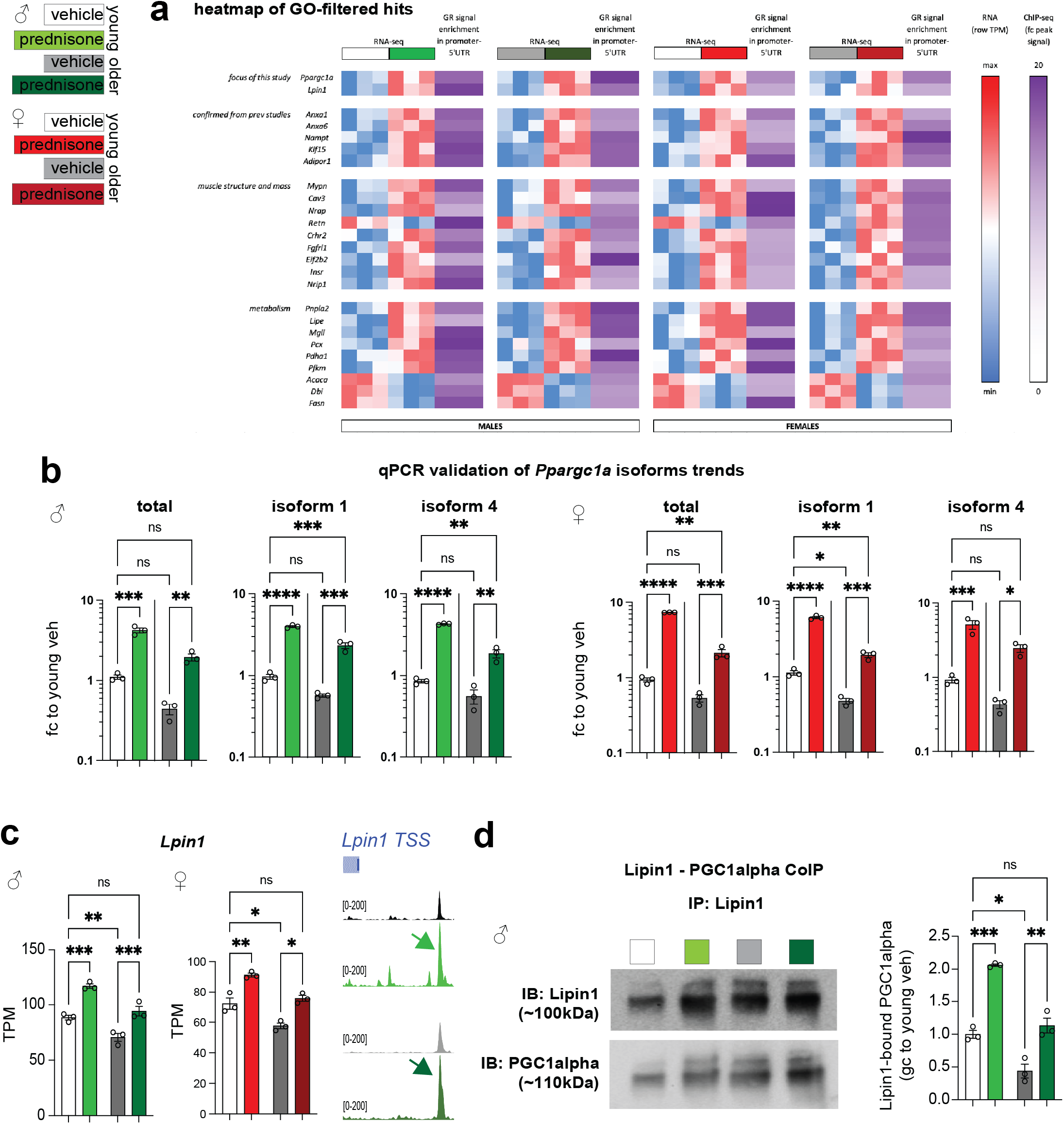
Related to Figure 2. Additional analyses regarding ChIP-seq X RNA-seq overlay and Ppargc1a-Lpin1 hits. **(a)** Heatmap summarizing RNA and GR signal trends for all GO-filtered hits according to sex/age groups. **(b)** qPCR analysis in quadriceps muscle RNA samples confirmed the RNA-seq trends in total Ppargc1a and isoforms 1/4 expression. **(c)** *Lpin1* expression was upregulated by treatment and rescued to young-like levels in older muscles in both males and females. Treatment increased the GR peak on *Lpin1* promoter in both age groups. **(d)** CoIP analysis from control and treated quadriceps muscles showed increased Lipin1-PGC1alpha interaction after treatment counteracting the aging-related effect. N=3/group; 2w ANOVA + Sidak: *, P<0.05; **, P<0.01; ***, P<0.001; ****, P<0.0001.

**Supplementary Figure 4.**
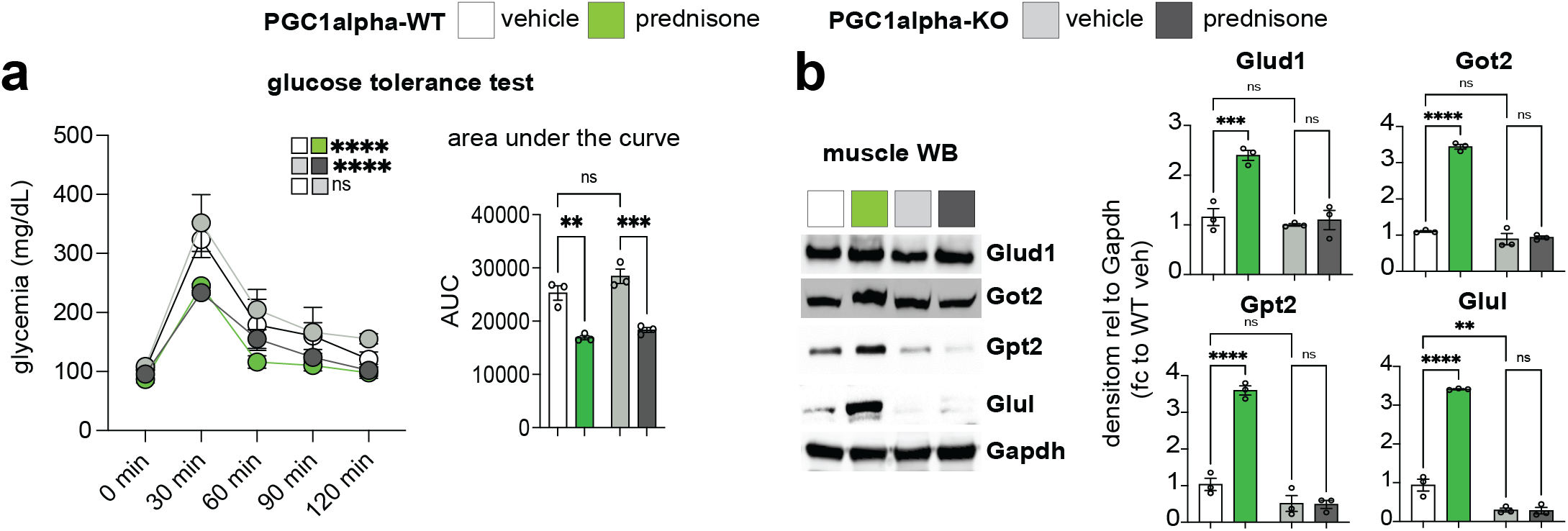
Related to Figures 3-4. Additional analyses with inducible PGC1alpha ablation in myofibers. **(a)** Treatment increased glucose tolerance regardless of myofiber PGC1alpha presence. **(b)** WB confirming the PGC1alpha-dependent upregulation of Glud1, Got2, Gpt2, Glul levels in muscle downstream of treatment. N=3/group; 2w ANOVA + Sidak: *, P<0.05; **, P<0.01; ***, P<0.001; ****, P<0.0001.

**Supplementary Figure 5.**
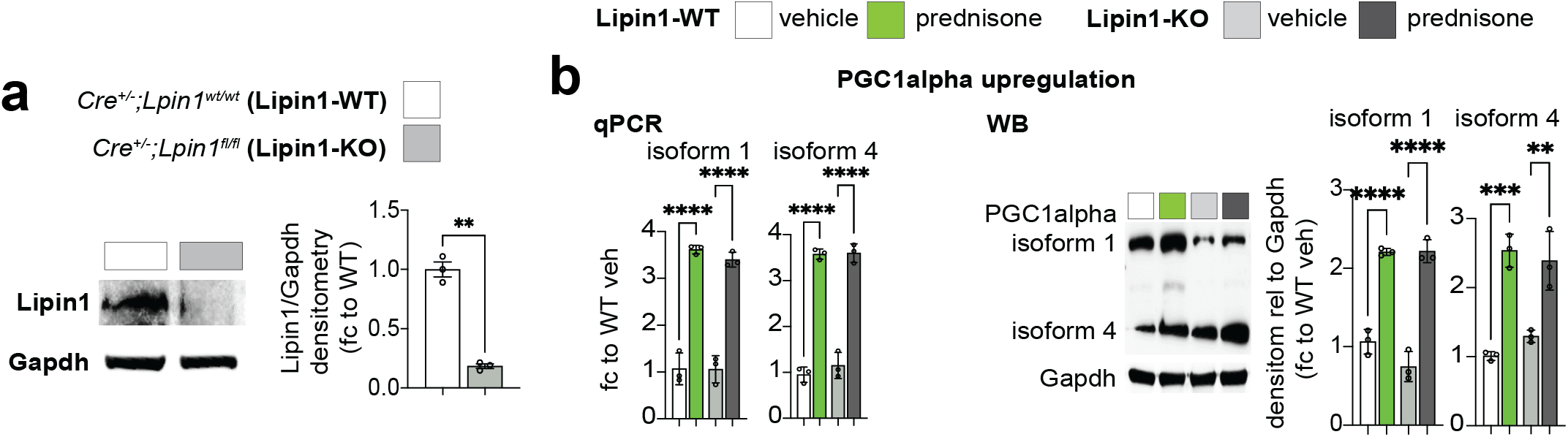
Related to Figure 5. Additional analyses with inducible Lipin1 ablation in myofibers. **(a)** Lipin1-KO validation. **(b)** Treatment increased expression of both isoforms 1 and 4 of PGC1alpha in muscle independently from myofiber Lipin1 presence. N=3/group; Welch’s t-test (a), 2w ANOVA + Sidak (b): *, P<0.05; **, P<0.01; ***, P<0.001; ****, P<0.0001.

## Notes

**Conflicts of interest –** MQ is listed as co-inventor on a patent application related to intermittent glucocorticoid use filed by Northwestern University (PCT/US2019/068618). All other authors declare no competing interests.

### Competing Interest Statement

MQ is listed as co-inventor on a patent application related to intermittent glucocorticoid use filed by Northwest-ern University (PCT/US2019/068618). All other authors declare no competing interests.

